# A Membrane-Targeted Photoswitch Potently Modulates Neuronal Firing

**DOI:** 10.1101/711077

**Authors:** Mattia L. DiFrancesco, Francesco Lodola, Elisabetta Colombo, Luca Maragliano, Giuseppe M. Paternò, Mattia Bramini, Simone Cimò, Letizia Colella, Daniele Fazzi, Cyril G. Eleftheriou, José Fernando Maya-Vetencourt, Chiara Bertarelli, Guglielmo Lanzani, Fabio Benfenati

**Affiliations:** Center for Synaptic Neuroscience, Istituto Italiano di Tecnologia, Largo Rosanna Benzi 10, 16132 Genova, Italy; IRCCS Ospedale Policlinico San Martino, Genova, Italy; Center for Nano Science and Technology, Istituto Italiano di Tecnologia, Via Pascoli 10, 20133, Milano, Italy; Dipartimento di Chimica, Materiali e Ingegneria Chimica *“Giulio Natta”*, Politecnico di Milano, Piazza L. da Vinci 32, 20133 Milano, Italy; Department of Chemistry, Institut für Physikalische Chemie, University of Cologne, Luxemburger Str. 116, D-50939 Köln, Germany

**Keywords:** photoexcitation, azobenzene derivatives, capacitance, lipid rafts

## Abstract

Optical technologies allowing modulation of neuronal activity at high spatio-temporal resolution are becoming paramount in neuroscience. We engineered novel light-sensitive molecules by adding polar groups to a hydrophobic backbone containing azobenzene and azepane moieties. We demonstrate that the probes stably partition into the plasma membrane, with affinity for lipid rafts, and cause thinning of the bilayer through their trans-dimerization in the dark. In neurons pulse-labeled with the compound, light induces a transient hyperpolarization followed by a delayed depolarization that triggers action potential firing. The fast hyperpolarization is attributable to a light-dependent decrease in capacitance due to membrane relaxation that follows disruption of the azobenzene dimers. The physiological effects are persistent and can be evoked in vivo after labeling the mouse somatosensory cortex. These data demonstrate the possibility to trigger neural activity in vitro and in vivo by modulating membrane capacitance, without directly affecting ion channels or local temperature.

---

Optical technologies for the modulation of neuronal activity are becoming increasingly important in cell biology and neuroscience (1, 2). Indeed, the possibility to obtain neuronal excitation or inhibition on demand not only has allowed an unprecedented power in interrogating and dissecting out the function of specific brain circuits, but has also opened new perspectives for treating neurological and psychiatric diseases, including blindness caused by photoreceptor degeneration (3).

Optogenetics is the pioneering technique in neuro-optical technologies. It consists in the expression of heterologous genes coding microbial opsins (light-dependent ion channels or transporters) that, acting as molecular sensor-actuators, transduce the light signal of appropriate wavelength into inward or outward ion current that excite or silence neurons, respectively (3, 4). However, optogenetics requires genetic modification of target cells that is a limitation for an easy translation to human therapy. Similar to optogenetics, the generation of tethered photoswitches based on azobenzene isomerization covalently linked to recombinant ligand-gated or voltage-gated ion channels has allowed modulating channel opening and closing in a light-dependent fashion (5). However, in addition to the need of expressing a recombinant channel, these photoswitches, such as maleimide-azobenzene-glutamate (MAG), were sensitive to the UV region of the spectrum that is harmful and poorly efficient in tissue penetration.

Recently, a new generation of azobenzene-based photoswitchable affinity labels has been demonstrated to photosensitize endogenous proteins without the requirement of genetic engineering (6–10). A first class of such probes in which the azobenzene group was flanked by a quaternary ammonium proved to be effective in blocking K^+^-channels and HCN-channels in response to UV light, thereby triggering neurons activation both in cultured neurons and in blind retinas injected intravitreally (11–13). From the original probe, new photoswitches sensitive to visible light and triggering light-reversible blockade of voltage- or ligand-gated ion channels has been generated (14–20). Most of these probes act intracellularly and relay on the exact positioning at the channel site.

Extracellular photostimulation by light-sensitive interfaces represent an alternative strategy. Extended planar organic interfaces were used to achieve light-dependent modulation of the electrical state of neurons (21–25) that, at high light intensities, also involved a thermal effect (23,26,27). Similar results were obtained by increasing the local temperature with IR illumination of absorbing materials in contact with cells (28, 29). The rise in temperature primarily increases the membrane capacitance that in turn depolarizes the target cell. This is likely due to a structural change of the charged membrane that becomes thinner by shortening/tilting the phospholipid tails with temperature and associated with an increase in the membrane area per phospholipid headgroup (30, 31). However, temperature rises of several degrees can be harmful to neurons, particularly if repeatedly administered.

In this work, we engineered an amphiphilic azobenzene photoswitch to obtain an intramembrane actuator for inducing *heatless* membrane stress/perturbation upon irradiation with visible light (32).

Our photochromic actuator contains either one (Ziapin1) or two (Ziapin2) polar heads that align with the phospholipid headgroups, and an azobenzene moiety end-capped with a hydrophobic azepane that can be folded/unfolded in a light-dependent manner. Incubation of the compounds with primary neurons showed that the molecules spontaneously partition into the membrane, where they preferentially distribute to membrane rafts. Light-dependent *trans➔cis* isomerization of the more active compound Ziapin2 caused a sharp and transient decrease in capacitance associated with hyperpolarization, which was followed by a sustained depolarization and firing of action potentials. Light-evoked stimulation of Ziapin2-labeled mouse somatosensory cortex activity was also observed *in vivo*. Molecular Dynamics (MD) simulations demonstrated that the *trans* isomers cause thinning of the bilayer through their trans-dimerization and that such membrane stress is relieved upon illumination following the *trans➔cis* isomerization that displaces the hydrophobic end-group from the membrane core.

## RESULTS

### Synthesis and spectroscopic characterization of the photochromic molecules

To synthetize light-sensitive amphiphilic probes targeting biological membranes and modulating their biophysical properties upon illumination, we engineered a hydrophobic backbone containing 4-4’ diaminoazobenzene group substituted on one side with an azepane and on the opposite side with alkyl chains that are ω-substituted with cationic groups, i.e. pyridinium or ammonium salts, using halides as counterion (i.e. Br^−^ or I^−^). The combination of the alkyl-substituted azobenzenes with a capping cation leads to amphiphilic species able to be driven inside the cell membrane. We synthesized two plasma membrane-targeted amphiphilic azobenzenes bearing either one (Ziapin1) or two (Ziapin2) pyridinium salts (**Fig. S1** and **Suppl. Materials**).

Azobenzene molecules undergo reversible *trans➔cis* isomerization upon illumination with visible radiation (**Fig. 1a**), with the reverse *cis➔trans* isomerization driven by either light or slower vibrational cooling owing to the thermodynamic stability of the *trans* isomer (typical energy difference ≈ 0.5 eV; 33-35). Ziapin2 shows the typical UV-vis absorption features of para amino-substituted azobenzenes, with a strong absorption peak centered at 470 nm (**Fig. 1b**) that can be attributed to the transitions of the *trans* isomer (35). Irradiation with blue light (450 nm) leads to *trans➔cis* isomerization that can be seen from the bleaching of the *trans* isomer absorption accompanied by the concomitant increase of the *cis* conformer absorption at 350-380 nm and 520-600 nm. The collective photoswitching dynamics upon light exposure, followed by monitoring the decrease of absorbance at 470 nm *vs*. irradiation time (**Fig. 1c**), reveals a well-defined and relatively fast photoreaction dynamics of Ziapin2 in DMSO, reaching a photostationary population after about 100 s of illumination and achieving a complete recovery with a t_1/2_ of 108 s in the dark at room temperature (36–38). The photoisomerisation process, which likely involves either the reduction of the N=N π-bond order followed by the twisting around the N-N bond or the inversion mechanism, occurs typically in the picosecond range, while the population dynamics toward the photostationary state takes much longer depending on the conformer thermodynamic stability and molecular environment (33, 36–40).

**Figure 1.**
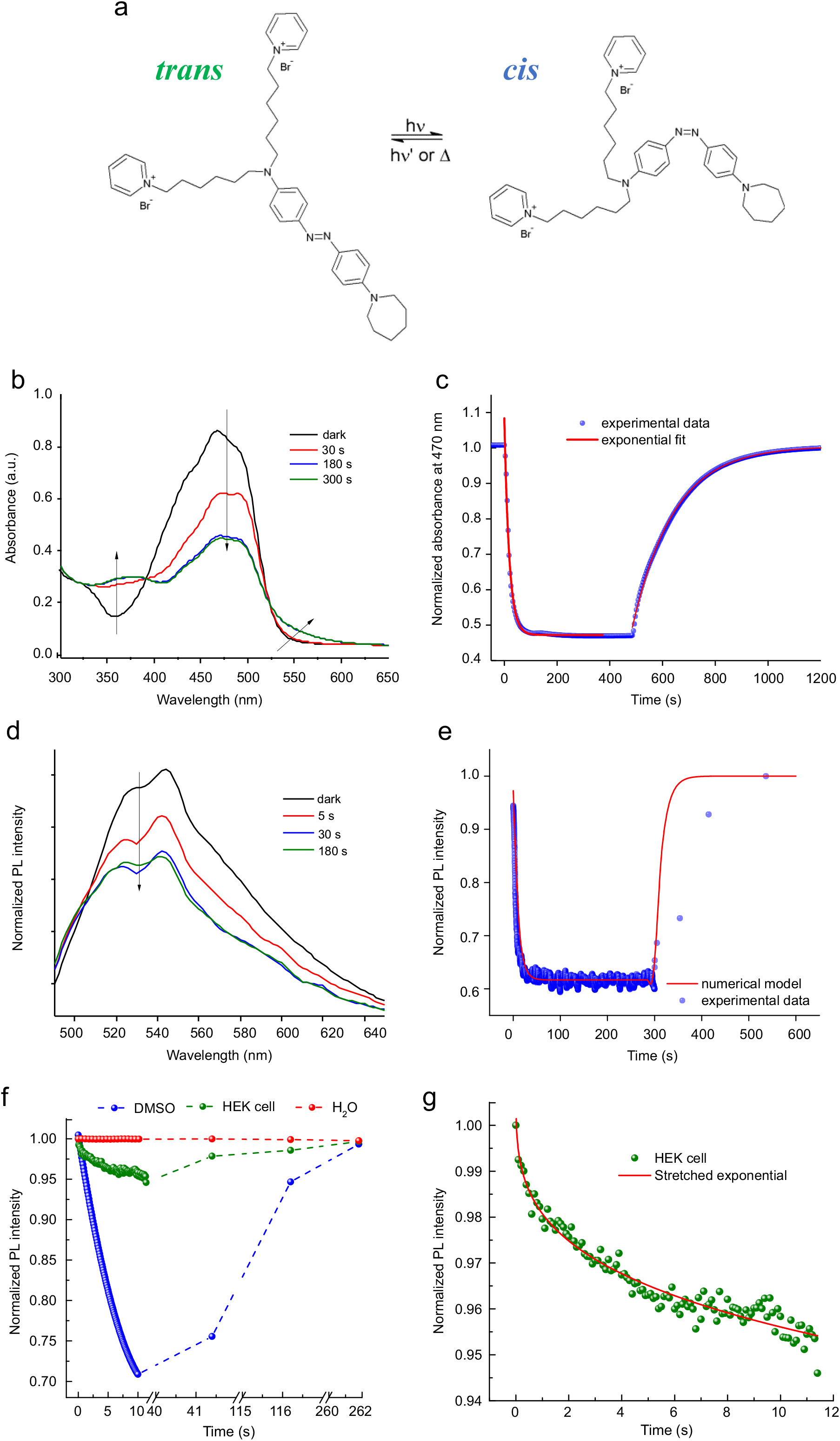
Spectroscopic characterization of *trans➔cis* isomerisation of Ziapin2 in various environments. **(a)** Schematics of the isomerisation process in Ziapin2. **(b,c)** Changes in the absorbance spectrum of Ziapin2 (25 μM in DMSO) as a function of time **(b)** and time-course of absorbance at 450 nm upon illumination with a diode laser **(c)**. **(d,e)** Changes in the photoluminescence (PL) spectrum of Ziapin2 (25 μM in DMSO) as a function of time **(d)** and time-course of the emission at 540 nm upon excitation at 450 nm with a Xenon lamp **(e)**. The red line is the numerical model employed to describe the time-evolution of the fluorescence signal. **(f)** Photoswitching/relaxation dynamics of Ziapin2 in DMSO, HEK293 cells and water acquired by exciting at 450 nm and collecting the emission at 540, 580 and 620 nm, respectively. **(g)** Zoom on the PL dynamics in HEK cells, highlighting the stretched exponential decay. Such function takes into account the distribution of relaxation times occurring in disordered environments.

The steep time-dependent decrease in Ziapin2 photoluminescence (PL) at 540 nm in DMSO upon blue light exposure (**Fig. 1d,e**) can be related to PL quenching of the *trans* conformer due the photoisomerization reaction and to the negligible PL quantum yield of the *cis* isomer. The excitation profile of the normalized change in PL upon illumination (Fig. S2a,b) follows the absorption profile of the *trans* isomer, confirming that it is indeed related to the isomerization process. By employing a numerical model (red line in Fig. 1e; see Online Methods) to describe the fluorescence changes *vs* the illumination time, we estimated a *trans➔cis* photoisomerization coefficient *kτ* = 16.9 cm^2^ J^−1^ and a *cis➔trans* thermalization rate *y* = 0.06 s^−1^. This validates Ziapin2 PL as valuable tool to monitor the photoswitching behavior of azobenzene.

The kinetics of Ziapin2 photoswitching was strongly dependent on the probe environment. Thus, a complete suppression of the Ziapin2 photoswitching was observed in aqueous environments as compared to DMSO (**Fig. 1f**), likely caused by strong aggregation (41). The formation of aggregates was also corroborated by the dramatic redshift (82 nm) of Ziapin2 PL (**Fig. S2c**). When Ziapin2 properties were assessed in the cell membrane environment, or in a micellar sodium dodecyl sulfate (SDS) solution that reproduces the lipid bilayer environment (42, 43), both spectral features and isomerization characteristics were recovered. The emission spectra of the Ziapin2-membrane and Ziapin2-SDS essentially coincide and lie between the spectra in DMSO and water (**Fig. S2c**).

Similarly, the photoswitching dynamics ability in either SDS micelles (**Fig. S2d**) or cell membranes (**Fig. 1f**) resulted to be intermediate between the fast photoswitching behavior in DMSO and the frozen photodynamics observed in aqueous media All these findings suggest that the partition of the molecule in lipid membranes avoids the aggregation of Ziapin2, enabling effective isomerization and light-controlled photoswitching.

The PL decay in cell membranes (**Fig. 1g**) was best fitted by a stretched exponential function (see Online Methods), describing the distribution of the relaxation times in disordered environments such as regions of the cell membrane with different viscosity features (e.g. lipid rafts) or local changes in the membrane status (i.e. phase or thickness). While the average distribution of relaxation times was centered at 33 s, an initial fast decay below 100 ms was present, consistent with the fast-biological response observed in electrophysiology experiments. It is worth noting that here we follow the evolution in time of the isomeric populations in the ensemble, whereas the molecular event of photoisomerization occurs in the sub-ps time scale. The single event can thus be extremely fast, but a sizable change is likely associated to a macroscopic fraction of isomerized molecules.

### Ziapin2 rapidly partitions to the neuronal plasma membrane and lipid rafts

To confirm the suggestions based on the spectroscopic characterization, we first simulated the spontaneous permeation of Ziapin2 in the model POPC membrane. We performed three distinct Molecular Dynamics (MD) simulations (two lasting 175 ns and one 300 ns) with randomized initial conditions after having placed one molecule of Ziapin2 in *trans* conformation in the water region, parallel to the membrane and in different orientations. In all three simulations, the molecule entered the bilayer after a time of about 40 ns, 80 ns and 100 ns, and remained in the membrane for the rest of the trajectories (**Fig. 2a,b**). We observed the same insertion mechanism in all three simulations: Ziapin2 entered the membrane very rapidly by first piercing it with the azobenzene side, and then moving towards the center of the bilayer by keeping the elongated axis almost parallel to the bilayer normal (**Fig. 2a**; I-IV). The insertion stopped when both the positively charged pyridine rings of Ziapin2 were at the level of the lipid heads, coordinated by negatively charged phosphate groups. The location along the bilayer normal was stationary for the rest of the simulations showing only limited fluctuations. Interestingly, in a different simulation, one Ziapin2 molecule placed at the center of the bilayer and parallel to it, reached very rapidly (< 5 ns) the same equilibrium position, parallel to the bilayer normal. The insertion process was studied quantitatively by calculating the free energy profile for moving Ziapin2 from bulk water into the bilayer (**Fig. 2c**). The free energy curve shows essentially no barrier for Ziapin2 adsorption in the membrane, a pronounced minimum at the same equilibrium position that was spontaneously found in the above simulations, and a barrier for membrane desorption to water of ≅ 12 kcal·mol^−1^ (0.52 eV).

**Figure 2.**
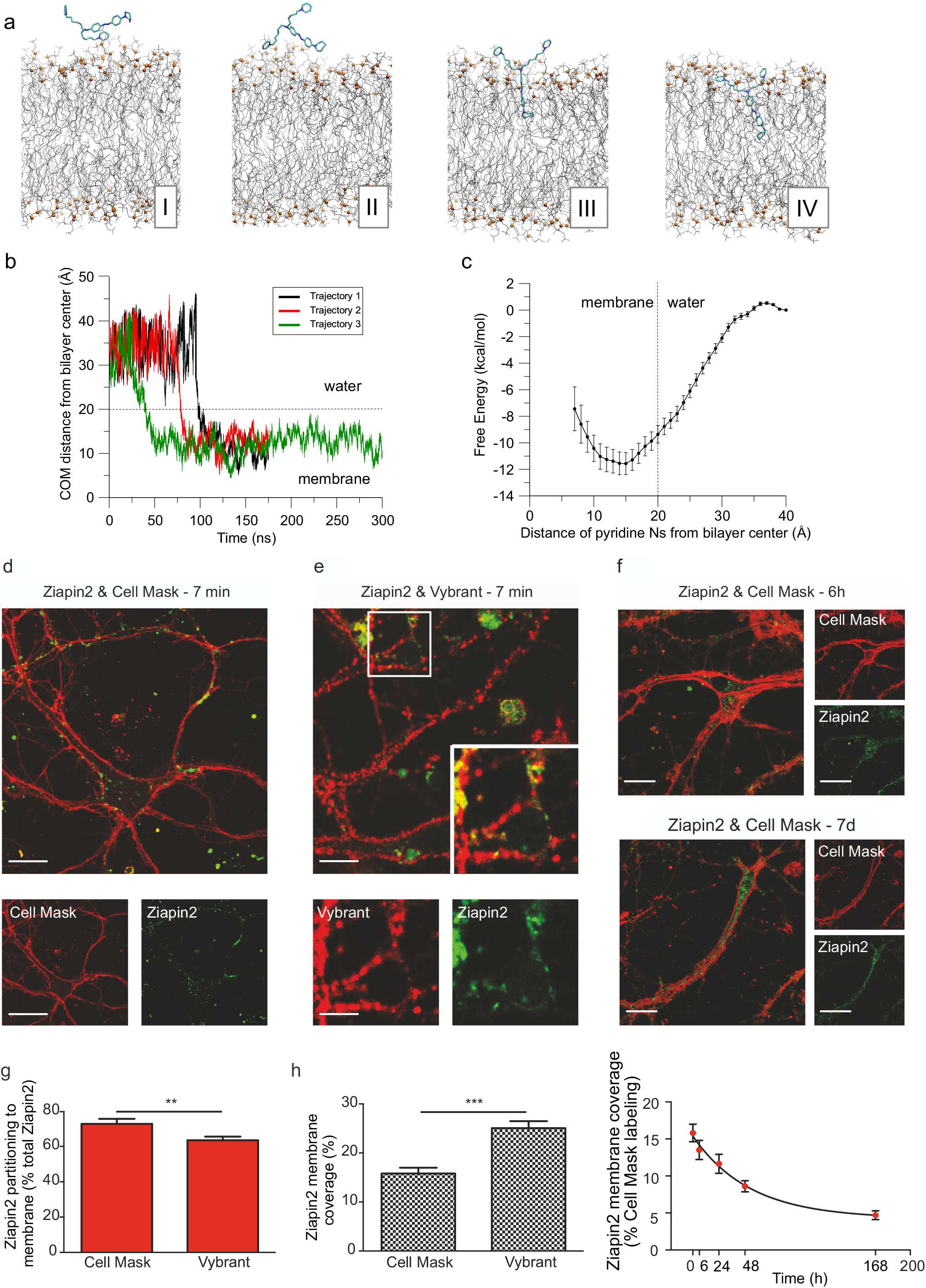
MD simulations in a plasma membrane model and targeting of Ziapin2 to the plasma membrane and lipid rafts in neurons. **(a)** Snapshots extracted from an MD simulation showing Ziapin2 *(trans)* spontaneously entering the membrane (POPC lipid model) at consecutive time frames (I-IV); lipid phosphate atoms are shown as orange spheres, and acyl chains as grey lines; water molecules are not reported for clarity. **(b)** Time dependence of the distance between the center of mass (COM) of Ziapin2 and the bilayer center in three independent simulations; the dashed line indicates the interface between water and lipid head groups. **(c)** Free Energy profile for Ziapin2 *(trans)* entering the membrane bilayer, calculated versus the distance between the bilayer center and the center of mass of the two pyridinic nitrogens of Ziapin2. **(d,e)** Primary neurons pulse exposed to Ziapin2 for 7 min were stained with specific membrane dyes to evaluate and quantify the co-localization of the molecule with the plasma membrane. Cell Mask staining was used to visualize the whole membrane of neurons (**d**, in red; Ziapin2 in green), while Vybrant™ Alexa Fluor™ 555 lipid rafts labeling kit was used for the identification of lipid rafts (**e**, in red; Ziapin2 in green). The Vibrant/Cell Mask surface ratio was 72 ± 4%. Scale bars: **(d)** 10 and 20 μm for large and small panels, respectively; (**e**) 10 and 4 μm for large and small panels, respectively. **(f)** The Cell Mask/Ziapin2 double staining is shown 6 h (upper panel) and 7 d (lower panel) after the initial exposure. Scale bars, 10 and 20 μm for large and small panels, respectively. **(g)** After Cell Mask/Ziapin2 and Vybrant/Ziapin2 double staining and z-stack confocal imaging, the partitioning of Ziapin2 to the plasma membrane and to lipid rafts was evaluated as the percentage of total cell Ziapin2 colocalizing with the respective membrane staining after 7 min of pulse incubation and subsequent washout (time 0). **(h)** Percentage of total plasma membrane (Cell Mask) or of lipid raft domains (Vybrant) positive for Ziapin2 after 7 min of pulse incubation (time 0; left panel). On the right, the percentage of plasma membrane (Cell Mask) positive for Ziapin2 is shown as a function of time up to 7 days after the initial exposure. The progressive decay was fitted using a one-component exponential decay function (t_1/2_ = 36.4 h).

To follow the subcellular localization of Ziapin2, primary hippocampal neurons were pulse loaded for 7 min, live-stained with Cell Mask after removal of the unbound molecule and subjected to 3D z-stack confocal imaging (**Fig. 2d**). The quantification revealed that the majority (>70%) of Ziapin2 was localized to the neuronal membrane where it covered the 15% of the total membrane surface (**Fig. 2g** and **2h**, left panel). The close overlap between Ziapin2 and the plasma membrane was also demonstrated by the ability of Ziapin2 emission to excite the live plasma membrane reporter Cell Mask (**Suppl. Materials and Fig. S3**). As the plasma membrane is in dynamic equilibrium with intracellular compartments through intense trafficking events, we analyzed the lifetime of plasma membrane Ziapin2 in primary neurons. Indeed, a slow and progressive decrease of plasma membrane coverage by Ziapin2 over time, characterized by a slow t_1/2_ of 36.4 h, was observed, (**Fig. 2h**, right panel), paralleled by an increased Ziapin2 fluorescence in intracellular compartments.

Lipid rafts are microdomains characterized by high concentration of cholesterol, glycosphingolipids and a variety of signaling and scaffolding proteins, including ion channels (44). We also evaluated the localization of Ziapin2 at lipid rafts by live-labeling with the cholera toxin B-subunit (Vybrant 555) that specifically recognizes the GM1 ganglioside (**Fig. 2e**). The percentage of Ziapin2 colocalizing with lipid rafts (~60%) was only slightly smaller than that observed with Cell Mask, with the molecule covering a much higher proportion (~25%) of the total raft surface (**Fig. 2g,h**; left panels). These findings suggest that Ziapin2 strongly partitions to the membrane bilayer with a preferential affinity for lipid rafts.

### Photoisomerization of Ziapin2 alters the membrane potential of primary hippocampal neurons

As Ziapin2 targets and isomerizes within the membrane and is predicted to affect membrane thickness, we investigated the effects of photoisomerization in primary hippocampal neurons labeled with 5 μM Ziapin2 and stimulated at 470 nm with a power density of 18 mW/mm^2^. To distinguish cell autonomous effects from the effects of the reverberant network of synaptic connections that characterizes primary neuronal cultures, neurons were recorded in the presence or absence of blockers of excitatory and inhibitory synaptic transmission (**Fig. 3b-d**).

**Figure 3.**
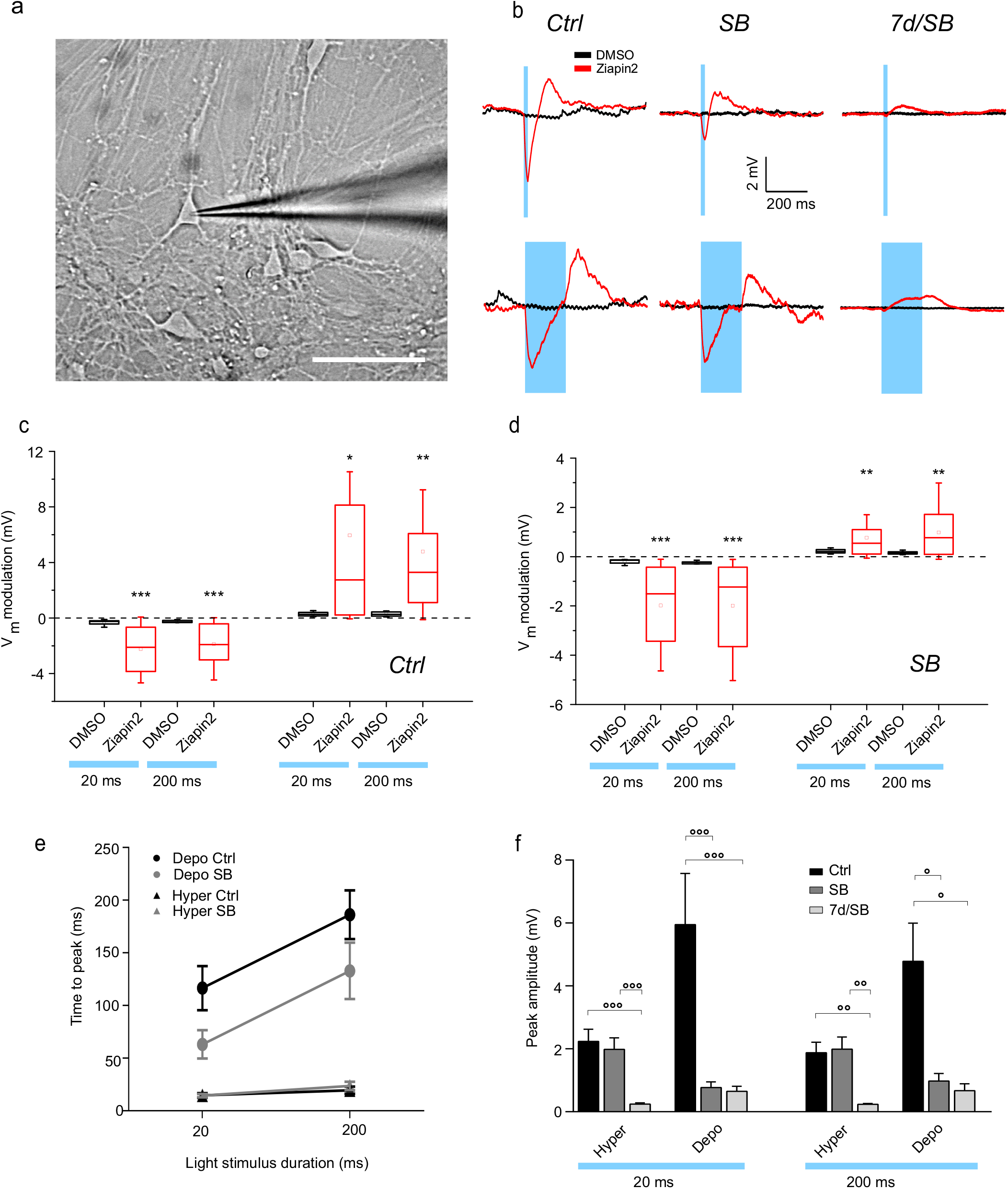
Light-evoked membrane voltage modulation in primary neurons by patch-clamp recordings. **(a)** Primary hippocampal neurons at 14 DIV were incubated with Ziapin2 for 7 min, washed and recorded by patch-clamp either immediately after pulse labeling or 7 days after. Scale bar, 20 μm. **(b)** Representative current-clamp traces recorded from neurons incubated with either 0.25% (v/v) DMSO (black traces) or 5 μM Ziapin2 in DMSO (red traces) in the absence (Ctrl) or presence of synaptic blockers (SB; see Materials and Methods), and after 7 days of incubation in the presence of synaptic blockers (7d/SB). The duration of the light stimulation (20 and 200 ms) is shown as a cyan-shaded area (470 nm; 18 mW/mm^2^). **(c,d)** Box plots of the peak hyperpolarization (left) and peak depolarization (right) changes in primary neurons exposed to DMSO/Ziapin2 and subjected to 20/200 ms light stimulation in the absence (c; Ctrl) or presence (d; SB) of synaptic blockers. Hyperpolarization and depolarization were measured as the minimum and maximum voltage, respectively, reached within 350 ms from light-onset. The box plots show that the peak hyperpolarization response generated by the presence of Ziapin2 is a cell autonomous response and is not affected by the presence of blockers of synaptic transmission, while depolarization, already present in synaptically isolated neurons, is enhanced by active synaptic transmission. **(e)** Time-to-peak hyperpolarization and depolarization as a function of the light stimulus duration **(f)** in Ctrl and SB conditions. Data (means ± sem) represent the time necessary to reach the minimum and maximum membrane voltages in the above-mentioned time windows. **(f)** Persistence of the depolarizing response over time. Peak amplitudes (means ± sem) of hyperpolarization and depolarization recorded immediately after pulse labeling and washout in the absence (Ctrl) or presence (SB) of synaptic blockers and 7 days after pulse labeling in the presence of synaptic blockers (7d/SB). All experiments with neurons were carried out at 24 ± 1 °C. * p<0.05; ** p<0.01; *** p<0.001; Ziapin2 *vs* DMSO, Mann Whitney *U*-test. ° p<0.05; °° p<0.01; °°° p<0.001; Ctrl *vs* SB *vs* 7d/SB, one-way ANOVA/Holm-Sidak test (Ziapin2 – Ctrl, SB, 7d/SB: N = 19, 20, 15 and N = 20, 19, 14 for 20 and 200 ms respectively; DMSO – Ctrl, SB, 7d/SB: N = 10, 7, 10).

Ziapin2-labeled hippocampal neurons showed a clear biphasic modulation of the membrane potential characterized by an early hyperpolarization, followed by a much larger depolarization (**Fig. 3b**). The effect was specific for Ziapin2, as no light-dependent effects could be described in neurons incubated with vehicle (DMSO; **Fig. 3b,d**). The amplitude of the hyperpolarization and depolarization peaks evoked by either 20 or 200 ms light stimulation were of similar magnitude and resulted highly significant with respect to the corresponding Vm oscillations induced by the same light stimuli in vehicle-treated neurons (**Fig. 3c,d**). While the hyperpolarization peak occurred with a relatively similar delay when stimulating for 20 or 200 ms, the peak depolarization was delayed with 200 ms stimuli (**Fig. 3e**). When the light-dependent effects were studied in the presence of synaptic blockers, a qualitatively similar and significant biphasic modulation of V_m_ was observed, confirming that the effects of illumination represent a cell autonomous effect (**Fig. 3b,c**). Interestingly, while synaptic blockade did not affect the magnitude of hyperpolarization or the timing of both hyperpolarization and depolarization responses, it significantly decreased the amplitude of the depolarization peak indicating that network excitatory transmission contributes to the late depolarization phase (**Fig. 3e,f**). To determine the light-induced current, we performed light-stimulation in whole-cell voltage-clamped Ziapin2-loaded neurons stimulated by 20 or 200 ms light pulses. Light stimulation elicited an outward current at negative potentials that became inward at strongly depolarized potentials, with an inversion between −60 and −30 mV (**Fig. S4**).

Morphological studies indicated that ~20% of the initial Ziapin2 labeling is still present on the plasma membrane seven days after the initial loading. Thus, we evaluated whether the residual Ziapin2 was still able to elicit the observed physiological responses. Indeed, still significant, albeit smaller, hyperpolarization and late depolarization responses were observed (**Fig. 3b,d**), suggesting a sustained effect of the photosensitization of the neuronal membrane, in spite of the membrane turnover progressively transferring the molecule to the intracellular compartments.

### Ziapin2 incorporation and photoisomerization within the plasma membrane induce capacitance changes by affecting membrane thickness

It has been shown that temperature-dependent changes in membrane capacitance can induce changes in membrane potential (26, 28). Thus, we performed simulations of Ziapin2 in both *trans* and *cis* conformations (200 ns/trajectory) in the membrane bilayer starting with the molecule in the equilibrium position. For the *cis* conformation, the initial position was determined by aligning the pyridine branches with those of the *trans* molecule. Our results show that in both cases the position along the bilayer normal is stationary over the whole trajectory, while the orientation fluctuates between a parallel and a perpendicular state. Spatially resolved membrane thickness maps calculated from simulations of multiple Ziapin2 molecules (4 and 8 Ziapin2 molecules per 160 lipids) revealed the presence of thinner regions in the bilayer when Ziapin2 molecules adopt the *trans* conformation than when it is in the *cis* state (**Fig. 4a,b**). We therefore investigated whether these thinning regions were due to particular spatial arrangements of the Ziapin2 molecules, observing that molecules in *trans* conformation anchored to the opposite leaflets of the membrane can in fact dimerize via backbone interaction. At the same time, the lipid heads orient themselves toward the center of the bilayer because of the electrostatic interaction between the pyridine rings with the phosphate head-groups, resulting in a local depression of the membrane (**Fig. 4c**). On the other hand, when Ziapin2 is in *cis* conformation, the azepane-substituted aniline of opposed molecules are too far from each other and do not dimerize, leaving the membrane thickness unperturbed (**Fig. 4d**). Accelerated MD simulations applied to a well-accepted raft model revealed that the same membrane push-pull effect also occurs in lipid rafts (**Fig. S5; Suppl. Materials**).

**Figure 4.**
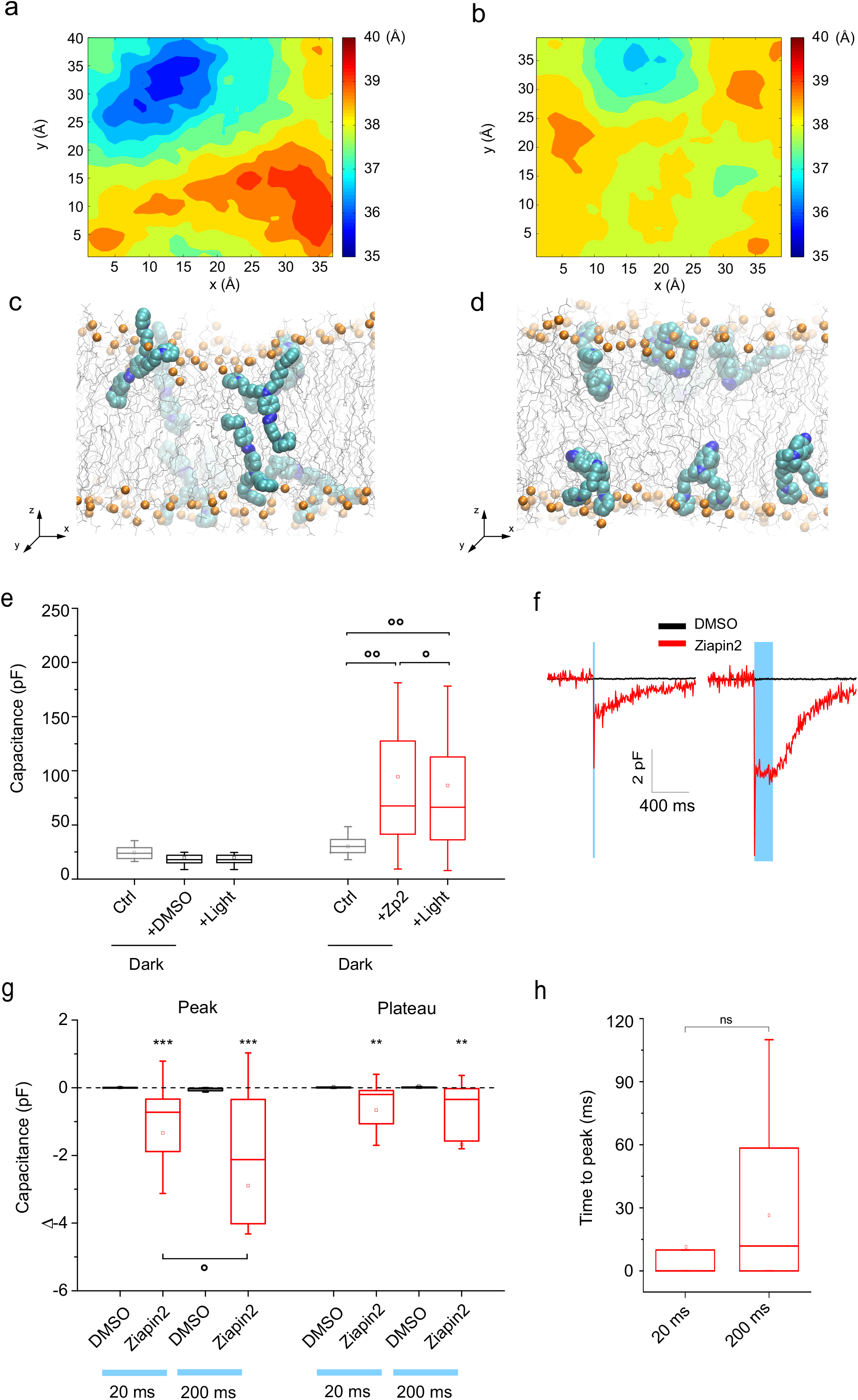
MD simulations of membrane thickness and experimental membrane capacitance changes in primary neurons loaded with Ziapin2 upon light stimulation. **(a,b)** Average membrane thickness maps, shown perpendicular to the bilayer plane, for the high-concentration (8 molecules) simulations of Ziapin2 in *trans* and *cis* conformation, respectively. Axis and thickness values are in Å. **(c,d)** Snapshots of from high-concentration simulations of Ziapin2 (8 molecules) in the *trans* and *cis* conformations, respectively. **(e)** Box plots of the cell capacitance changes after exposure of neurons to either DMSO (0.25% v/v; black) or Ziapin2 (5 μM in DMSO; red) in the dark (DMSO N=18; Ziapin2 N=24) and subsequent illumination for 200 ms in the presence of the compound and of synaptic blockers. **(f)** Representative averaged capacitance traces of neurons pulse labeled with either DMSO (0.25% v/v; black traces) or Ziapin2 (5 μM in DMSO; red traces), washed and recorded in current-clamp configuration in the presence of synaptic blockers before and after light stimulation with either 20 ms (left panel) or 200 ms (right panel) light pulses (470 nm; 18 mW/mm^2^; cyan-shaded areas). **(g)** Box plots of the peak capacitance changes (left: peak change; right: plateau change) evoked in neurons exposed to DMSO/Ziapin2 by 20 and 200 ms light stimuli (N = 9, 24, 9, 22 for the four experimental groups, respectively). **(h)** Time-to-peak capacitance change evoked by 20 and 200 ms light stimulation is reported (N= 24 per each experimental group). The dynamics of the capacitance changes were independent of the duration of the light stimulus. * p<0.05; ** p<0.01; Student’s t-test (e); ** p<0.01; *** p<0.001; Ziapin2 *vs* DMSO. Mann Whitney’s *U*-test (g); ° p<0.05, 200 *vs* 20 ms, one-way ANOVA/Holm-Sidak’s tests (g).

Based on these results, we experimentally checked whether loading primary neurons with Ziapin2 in the dark *(trans* configuration) in the presence of synaptic blockers had any effect on cell capacitance (**Fig. 4e**). While the basal values were homogeneous (24.9 ± 4.1 pF), indicating a reproducible size of primary neurons, the addition of Ziapin2 induced a marked increase in capacitance (up to 75.6 ± 22.8 pF) attributable to bilayer thinning caused by Ziapin2 homo-dimerization in *trans* and/or changes in the membrane dielectric constant (see Suppl. Materials). Under these conditions, the light-induced *trans➔cis* photoconversion of Ziapin2 led to a significant, albeit smaller, decrease of capacitance towards the basal values (65.8 ± 17.8 pF; **Fig. 4f**) likely due to a local membrane relaxation to the natural thickness. The fact that illumination was unable to bring back the capacitance to the basal level observed before addition of the compound can be explained by the observation that only a fraction of the *trans* molecules photoisomerize, and that photoisomerization *per se* may not affect the dielectric constant.

When measurements were performed after pulse labeling and removal of the Ziapin2-containing medium, capacitance fluctuations were characterized by a fast decrease that peaked at the light stimulus onset followed by a light-on steady-state level proportional to the stimulus duration, and a slow return to the pre-illumination level after stimulus offset (**Fig. 4f**). Both peak and offset changes in capacitance evoked by light showed a significant decrease with respect to vehicle-treated neurons (**Fig. 4g**). In the vast majority of neurons, the peak change in capacitance was almost instantaneous, and the delay between the light onset and the capacitance peak was not significantly different between 20 and 200 ms of light stimulation (**Fig. 4h**). The strict temporal correlation between the hyperpolarizing outward current and the change in membrane capacitance indicate that the light-induced membrane relaxation and the resulting increase in membrane thickness are responsible for the physiological effects (see **Suppl. Materials**).

### Primary hippocampal neurons exposed to Ziapin2 exhibit light-dependent firing

The described opto-mechanical effect on the membrane capacitance and voltage opens the possibility of a light-driven modulation of neuronal firing activity. To test this hypothesis, we evaluated the ability of light stimulation to elicit action potential (AP) firing in hippocampal neurons loaded with Ziapin2 shortly after and seven days after the initial pulse loading (**Fig. 5a-c**). Light stimulation elicited a significant increase of AP frequency in the absence of synaptic blockers (**Fig. 5a**). Such result was even more striking when neurons were measured in the presence of synaptic blockers that virtually abolished spontaneous light-independent firing (**Fig. 5b,c**). Peristimulus time histogram (PSTH) analysis showed that light reliably and significantly induced AP firing activity both in the absence and presence of synaptic blockers. In both cases, the strong increase in AP firing peaked after the light offset for short (20 ms) stimuli and during the late phase of illumination in the case of long (200 ms) stimuli (**Fig. 5d,e**), exhibits a strict time-dependent correlation with the late depolarization phase (**Fig. 5g**) that reflects a modulatory mechanism acting at the level of single neurons. To determine the persistence of the light-response over time after the initial pulse labeling, we tested whether the modulation of neuronal activity could still be detected one week after Ziapin2 labeling. In spite of the observed decrease in Ziapin2 content at the neuronal membrane (see **Fig. 2**), we found a persistent modulation of AP firing in response to light (**Fig. 5c,f,h**). Interestingly, repetitive firing was obtained with a 30 s train stimulation of 200 ms light pulses at 1 Hz or with 20 ms light pulses at 5 Hz, with only occasional failures in the firing response (**Fig. 5i**).

**Figure 5.**
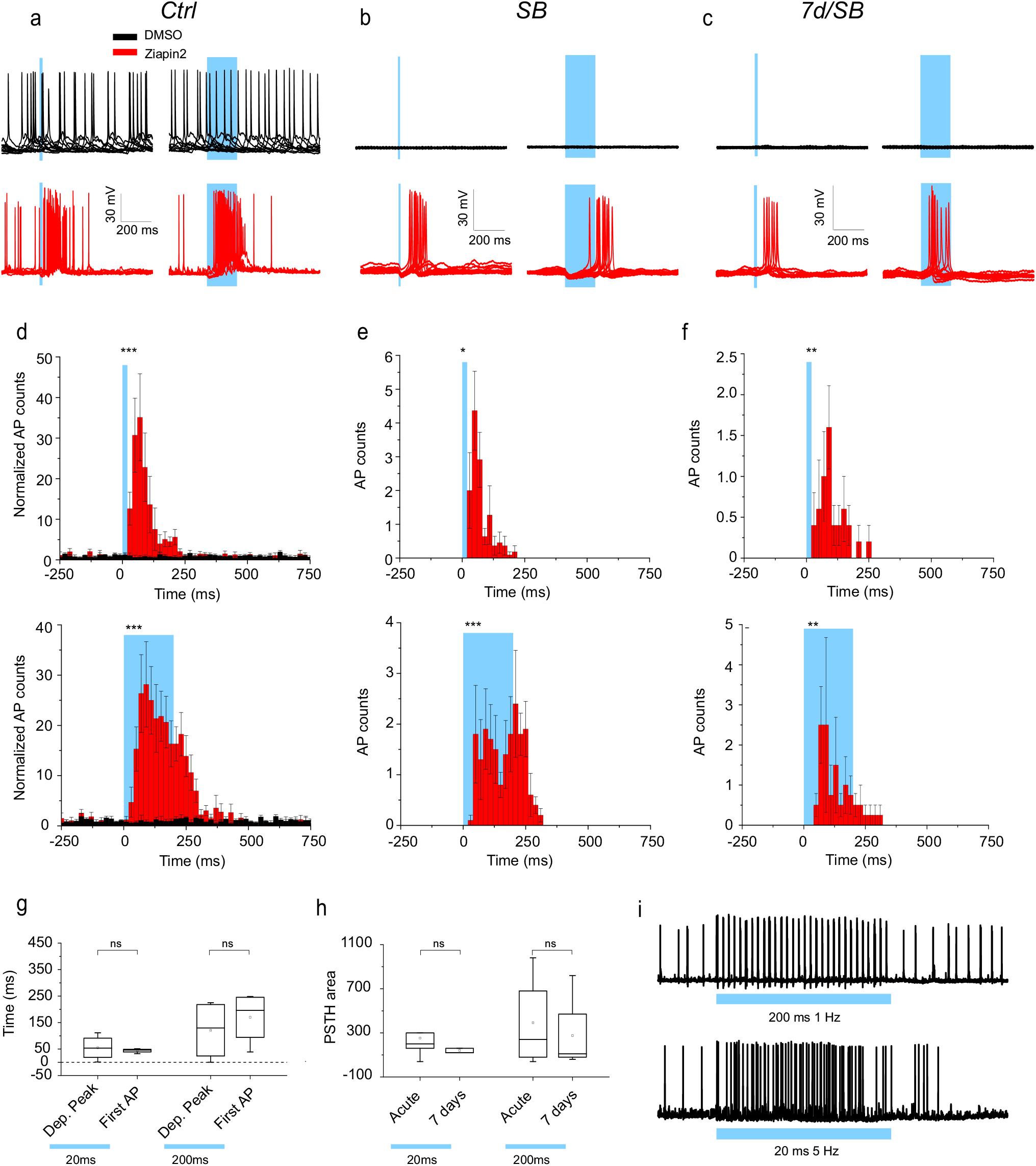
Light-evoked modulation of firing activity in primary neurons loaded with Ziapin2. **(a-c)** Representative averaged traces recorded in current-clamp configuration from neurons incubated with either DMSO (0.25% v/v, black traces) or Ziapin2 (5 μM in DMSO, red traces) in the absence (a; Ctrl) or presence (**b**; SB) of synaptic blockers. In (c) recordings were performed 7 days after Ziapin2 pulse labeling (7d/SB). The light stimulation (470 nm; 18 mW/mm^2^) is shown as a cyan-shaded area. (**d-f**) Peristimulus time histograms (PSTHs; bin=20 ms) reconstructed from the firing rate of neurons recorded in the absence (d) or presence of synaptic blockers 7 min (**e**) and 7 days (**f**) after DMSO/Ziapin2 pulse labeling, respectively and subjected to either 20 ms (upper panel) or 200 ms (lower panel) light stimulation (Ziapin2 – Ctrl, SB, 7d/SB: N = 7, 11, 5 and N = 6, 10, 4 for 20 and 200 ms respectively; DMSO – Ctrl, SB, 7d/SB: N = 4, 7, 10 for both durations). *** p<0.001 DMSO *vs* Ziapin2, Mann-Whitney Otest on 160 ms bins (20 ms stimulation) and 240 ms bins (200 ms stimulation). (**g**) Time delays of the depolarization peak and of the first action potential (AP) event from the light-onset in neurons pulse labeled with Ziapin2 (5 μM) in the presence of synaptic blockers and photostimulated for either 20 or 200 ms. Data were obtained below and above threshold are from the same neurons. Wilcoxon matched-pairs signed rank test. (**h**) PSTH areas of AP firing in response to 20/200 ms light stimulation in the presence of synaptic blockers recorded either acutely or 7 days after Ziapin2 exposure. N=10 (acute); N=5 (7 days); Mann Whitney *U*-test. (**i**) Representative AP firing activities recorded from neurons incubated with Ziapin2 (5 μM) in the absence of synaptic blockers and stimulated with 200 ms light pulses administered at 1 Hz (upper traces) or with 20 ms light pulses administered at 5 Hz (lower traces).

### The membrane-anchoring module is an important determinant of the physiological effects of Ziapins

To verify the importance of surface anchoring for the membrane localization and light-dependent modulation of the passive and active membrane properties, we also analyzed Ziapin 1 which is characterized by only one ω-pyridinium substituted alkyl chain (**Fig. S1**) and sharing similar spectroscopic properties with Ziapin2 (**Fig. S6a,b**). The membrane thinning induced by interacting opposite molecules in *trans* conformation and reversible upon isomerization was also observed by simulating Ziapin 1 molecules in the same membrane model environment (**Fig. S6c,d**). However, as expected by the single ionic anchor, Ziapin1 partition into the membrane bilayer was less stable (**Fig. S7a-c**), with lower percentages of the total cell probe co-distributing with the plasma membrane and lower membrane coverage than those observed with Ziapin2 (**Fig. S7d,e**). This was also testified by the faster decrease of Ziapin 1 plasma membrane labeling over time, paralleled by an early appearance of Ziapin 1 fluorescence in intracellular compartments (**Fig. S7e**, right panel) and the lower degree of excitation of Cell Mask after Ziapin 1 excitation (**Fig. S7f-h**). Consistent with these data, the light-dependent modulation of membrane capacitance and membrane potential by Ziapin 1 in primary neurons was qualitatively similar, but less intense than that obtained with equal concentrations of Ziapin2 (**Fig. S8a-c**). Neurons loaded with Ziapin 1 also displayed light-induced AP firing, although the firing probability was significantly smaller than that of Ziapin2 and the effect on firing was not detectable one week after the initial exposure to the compound (**Fig. S8d,e**).

### *In vivo* labeling of cortical neurons with Ziapin2 induces light-evoked cortical responses

To investigate the possible light-dependent modulation of neural activity also *in vivo*, Ziapin2 (200 μM in 1 μl 10% DMSO) was stereotaxically injected over 20 min in the somatosensory cortex of mice that were subsequently implanted with a multielectrode array coupled with an optical fiber for photostimulation (**Fig. 6a**). When unfixed brain slices corresponding to the site of injection were analyzed for the intrinsic Ziapin2 fluorescence, the area of diffusion and uptake of the molecule by cortical cells was in the range of 1 mm in diameter (**Fig. 6b**). Optical stimulation for 20 and 200 ms with a 472 nm laser at various power densities induced a significant activation of cortical activity evaluated as extracellular local field potentials (LFPs) that was not present in vehicle-injected animals (**Fig. 6c**). In most electrodes, the light stimulus induced a prompt potential artifact that was followed by a field response that peaked at about 200 ms after the stimulus. Quantitative analysis of the amplitude of LFPs larger than 5-fold the standard deviation revealed that Ziapin2 induced a significant light power-dependent increase in the LFP peak amplitude with respect to vehicle-injected animals that was more pronounced in response to 200 ms stimuli (**Fig. 6d**).

**Fig. 6.**
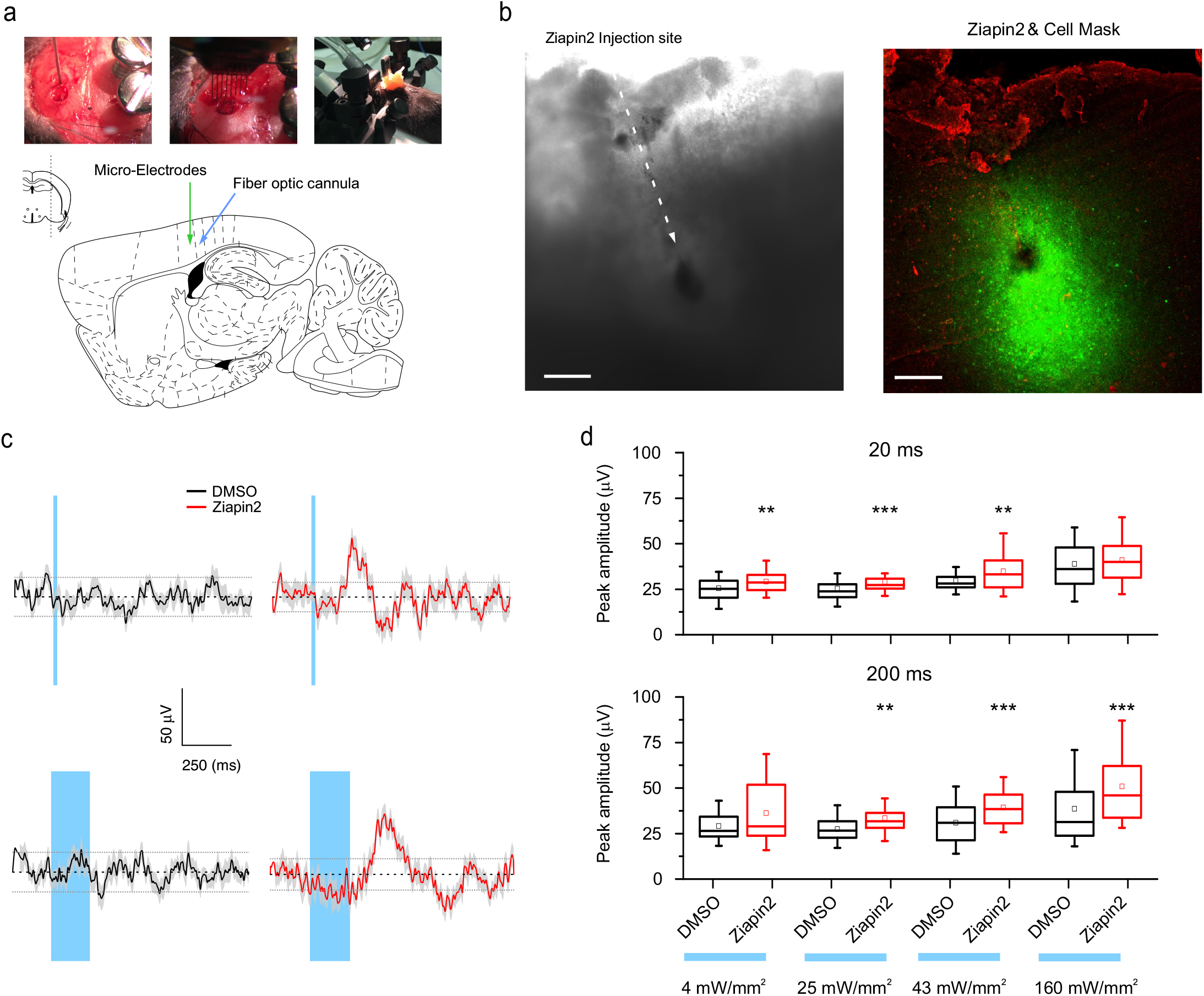
Light-evoked cortical responses *in vivo* in mice loaded with Ziapin2 in the somatosensory cortex. **(a)** Schematic representation of the stereotaxic injection of Ziapin2 (200 μM in 1 μl 10% DMSO) in the somatosensory cortex (S1ShNc, 2 mm anterior to lambda, 2 mm lateral to midline, and – 723 μm ventral to brain surface) and of the 16-microelectrode array implant coupled with optical fiber for photostimulation and local field potentials (LFPs) recordings. **(b)** Bright field image (left) and endogenous LC339 fluorescence micrograph (right) in an unfixed slice from the injected somatosensory cortex. The injection site and the diffusion of the compound are visible. In the right panel, the slice was counterstained with Cell Mask. Scale bar, 150 μm. **(c)** Representative recordings of LFPs evoked in the somatosensory cortex by 20 and 200 ms light stimulation (43 mW/mm^2^) in mice injected with either DMSO (black trace) or LC339 (red trace). The cyan-shaded areas represent the light stimulation. Potentials were considered significant above 2-fold the standard deviation range (broken horizontal lines). **(d)** Dose-response analysis of the LFPs in DMSO (black) or LC339 (red) injected mice as a function of the photostimulation power and duration (20 and 200 ms; top and bottom respectively). Stimulations at 25 and 43 mW/mm^2^ trigger a response in LC339 injected animals that is significantly different from DMSO treated animals, and at 4 mW/mm^2^ in the case of a 20 ms illumination. Higher power densities elicit a response that, possibly partially thermally driven, is significantly higher in presence of LC339. No significant changes with respect to DMSO alone were observed in the absence of the light stimulus and with the highest power used (160 mW/mm^2^). * p<0.05; ** p<0.01; *** p<0.001; Mann Whitney U-test (N= 3 mice for both experimental groups).

## DISCUSSION

We report here on a new opto-mechanical effect driven by simple intramembrane molecular machines composed of clustered photochromic molecules. MD simulations support the experimental observations showing that the hydrophobic azepane-substituted aniline in the amphiphilic azobenzenes on the two sides of the membrane interact when in the *trans* isomer, while they retract after photoconversion to the *cis* conformation. This brings about shrinkage of the membrane upon loading of Ziapin2 that relax to the natural membrane thickness consequent to photostimulation. While the membrane modification induced by the compound in the dark could be due to a change in local membrane thickness and/or dielectric constant, the dislocation of the non-polar moiety upon light exposure should not lead to sizable dielectric effects. This suggests that we are indeed exploiting a nanomachine inside the membrane that has a mechanical effect at the molecular scale.

The light-induced membrane relaxation increases membrane thickness, thus decreasing its capacitance. This change, in turn, is responsible for the hyperpolarization of the membrane potential, whose amplitude depends on the extent of membrane coverage and the dynamics of capacitance change. Several papers reported a link between a temperature-dependent decrease in membrane thickness, increased capacitance and membrane depolarization (26, 28–31).

The light-induced hyperpolarization is followed by a rebound depolarization wave that triggers action potential firing. We previously showed, using conjugated polymer interfaces (23, 27), that light-induced inhibition is followed by a rebound depolarization and firing, indicating that either the return of capacitance to the pre-stimulus levels or the activation of hyperpolarization-gated inward currents is responsible for neuronal activation. The depolarization amplitude and subsequent firing are enhanced by the excitatory synaptic transmission indicating that, although depolarization is a cell-autonomous effect it can generate a positive feedback within the network that amplifies light-evoked neuronal excitation. Interestingly, such neuronal activation was persistent *in vitro*, in spite of the decrease in the presence of the probe in the plasma membrane due to membrane turnover. Moreover, the photochromic molecule was effective in inducing a light-dependent electrical activation of the cortical networks after *in vivo* administration in the somatosensory cortex of the mouse, paving the way to their use in the therapy of degenerative diseases of the retina, as well as, of central nervous system.

In conclusion, our new amphiphilic photochromic molecules have several characteristics that differentiate them from previous compounds, namely: (i) marked affinity for the hydrophobic environment of the membrane; (ii) high tolerability and sensitivity to the visible spectrum; (iii) reversible induction local membrane deformations upon illumination that alter membrane capacitance, potential and firing in the absence of heat generation; (iv) effectiveness *in vivo*. In view of these features, these molecules display a high potential for future applications in neurosciences and biomedicine.

## Acknowledgements

The work was supported by the Italian Ministry of Health (project RF-2013-02358313 to GP, GL and FB) and Istituto Italiano di Tecnologia (pre-startup project to GL and FB). The support of Ra.Mo. Foundation (Milano, Italy), Fondazione 13 Marzo (Parma, Italy) and Rare Partners srl (Milano, Italy) are also acknowledged.

## Author contributions

C.B. designed the photoswitches and engineered the Ziapin compounds. S.C. and L.C. performed the synthesis of the Ziapin compounds. D.F. calculated the atomic charges and optimized coordinates. G.M.P. performed the spectroscopic characterization. L.M. performed molecular dynamics simulations. M.B. and E.C. studied the in vitro distribution of the Ziapin compounds in neurons. M.L.D., E.C. and F.L. performed the in vitro patch-clamp experiments and analyzed the data. J.F. M-V., E.C. and C.G.E. performed and analyzed the in vivo experiments. M.L.D., E.C., G.M.P. and F.L. contributed to paper writing. G.L., C.B. and F.B. conceived the work, G.L. and F.B. planned the experiments, analyzed the data and wrote the manuscript.

## Competing interests

The authors declare no competing interests.

## Additional information

Supplementary information is available for this paper.

## ONLINE METHODS

### Synthesis of the photochromic molecules

Unless otherwise stated, all chemicals and solvent were commercially available and used without further purification. Reactions of air- and water-sensitive reagents and intermediates were carried out in dried glassware and under argon atmosphere. If necessary, solvents were dried by means of conventional method and stored under argon. Thin layer chromatography (TLC) was performed by using silica gel on aluminum foil (Sigma Aldrich). NMR spectra were collected with a Bruker ARX400. Mass spectroscopy was carried out with a Bruker Esquire 3000 plus. 4,4’-diaminoazobenzene (1) was synthesized according to the previously reported procedure (45).

*4,4’-Bis-(N,N-di-w-bromohexyl)diaminoazobenzene (Azo-Br4), Azo-Br1, Azo-Br2*. 1.0 g of 1 is stirred in 130 ml of previously degassed acetonitrile. 2.60 g of K_2_CO_3_ and 7.5 ml of 1,6-dibromohexane are added portionwise to the reaction mixture and refluxed for 120 hours, while monitored by TLC. The reaction mixture is filtrated to and the solid is washed three times with diethyl ether, ethylacetate and dichloromethane. The excess of dibromohexane is removed under reduced pressure (3 10^−1^ mbar) at 60 °C. The raw material is purified by flash chromatography with silica gel to give 30 mg of the desired product. From the flash chromatography, 100 mg of *Azo-Br1* and *Azo-Br2* are also recovered. *Azo-Pyr1* (*Ziapin1*). 12 mg of Azo-Br1 are dissolved in 3 ml of pyridine and stirred at room temperature for 42 hrs. Then 3 ml of methanol are added and further stirred for 60 hrs. The excess of pyridine and methanol are removed from the reaction mixture under reduced pressure to give a solid that is washed with small portions of hexane.*Azo-Pyr2 (Ziapin2)*. 12 mg of Azo-Br2 are dissolved in 3 ml of pyridine and stirred at room temperature for 42 hrs. Then 3 ml of methanol are added and further stirred for 60 hrs. The excess of pyridine and methanol are removed from the reaction mixture under reduced pressure to give a solid that is washed with small portions of hexane.

### UV-VIS absorption measurements

For the UV-VIS absorption measurements, we used a Perkin Elmer Lambda 1050 spectrophotometer, equipped with deuterium (180-320 nm) and tungsten (320-3300 nm) lamps and three detectors (photomultiplier 180-860 nm, InGaAs 860-1300 nm and PbS 1300-3300 nm). All the absorption spectra were corrected for the reference spectra taken at 100% transmission (without the sample) at 0% transmission (with an internal attenuator), and for the background spectrum (DMSO only). To induce the isomerization process, we employed a continuous wave (CW) diode laser (OXXIUS) at 450 nm (incident power 25 mW). We placed the illumination source at 90° with the respect of the white light and detector. To acquire the isomerization collective kinetic of the azobenzene molecule in DMSO, we monitored the absorbance of the solution (DMSO 25 μM, 1 cm optical path) at 490 nm under illumination and during thermal relaxation in the dark.

### Photoluminescence measurements in solution

The PL measurements in solution (25 μM in DMSO, water and sodium dodecyl sulfate (SDS; 100 mM) were taken with a Horiba Nanolog Fluorimeter, equipped with a xenon lamp, two monochromators and two detectors (photomultiplier and InGaAs). To measure the time-evolution of the PL signal under continuous illumination, we excited at the maximum of the absorption peak (470 nm for the solution in DMSO and SDS, and 500 nm for the solution in water) and collected the emission at the peak maximum (540 nm, 580 nm and 620 nm for the solution in DMSO, water and SDS, respectively). To measure the PL signal during the *cis* to *trans* thermal relaxation, we collected the emission every 100 ms over 1 s of light exposure. The discrete points presented are the average of 10 scans collected over 1 s (the error bars represent the standard deviation). All the spectra were normalized to the lamp intensity.

### Numerical model of the PL signal dynamics in solution

The photoisomerization kinetics of azobenzene molecules is a well-known process that is widely described in the literature. Under illumination, the photoisomerization reaction takes place and a number of molecules in the thermodynamically stable *trans* state (*n_tra_*(*t*)) pass to the metastable *cis* state (*n_cis_*(*t*)). Such a process is described by the differential equation (1):

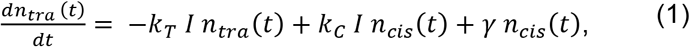

where *γ* is the rate of the *cis➔trans* thermal relaxation, *k_T_* and *k_C_* are the intensity normalized rate constants for the *trans➔cis* and *cis➔trans* photoisomerization processes, respectively.

The fraction of molecules in the *trans* and *cis* states can be defined as 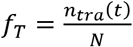 and 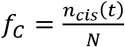, where *N* is the total number of molecules.

It is now possible to rewrite equation (1) as:

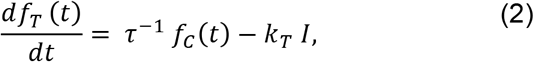

where 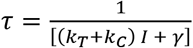 is the characteristic time-constant for the photoisomerization process. We solved such differential equation (2) by implementing the Euler method, in which a differential function can be written as an incremental ratio 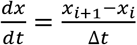 where *x_i_* is the value at the time *i* and *x*_*i*+1_ is the value at the time i+1. Therefore, to obtain the numerical solution, we expressed the differential equation in terms of the incremental ratio and we calculated the value of *f_T_* at the time i+1. We employed a custom Matlab script to implement this method.

### Photoluminescence (PL) measurements in membranes

Human Embryo Kidney cells (HEK293), used as a source of cell membranes, were obtained from ATCC. The emission of azobenzene in the membrane environment at 450 nm was performed by using a CW diode laser (excitation energy of 10 mW mm^−2^, matching the one of electrophysiology experiments). The emission was collected with a 50x objective (Zeiss), filtered to remove the wavelength excitation and sent to the camera (Hamamatsu, acquisition time 100 ms). The system was illuminated for a shorter time (10 s) than the PL measurements in solution, to avoid cells damaging. The relaxation dynamic was studied by employing the same method described above for the measurements in solution. The decay dynamics in cells was fitted via a stretched exponential function *y* = *A*· exp (-*t/τ*)^β^, where *A* is a pre-exponential constant, ***τ*** is the average relaxation time and *β* is shape parameter (0 < β ≤ 1) that measures the deviation from a simple exponential decay.

### Molecular dynamics (MD) simulations

All-atom Molecular Dynamics (MD) simulations of Ziapin2 in *trans* and *cis* conformations in a model membrane bilayer of 1-palmitoyl-2-oleoyl-phosphatidylcholine (POPC) made of 160 lipid molecules (80 per leaflet) and water. A starting conformation for the bilayer in water was generated using pre-equilibrated lipid structures from the CHARMM-GUI webserver (46). All simulations were run with NAMD v2.12 code (47) using the CHARMM36 force field (48) and TIP3P model for water molecules. CHARMM-compatible topology and parameters for the ZIAPIN1 and ZIAPIN2 molecules were obtained using the CHARMM General Force Field (CGenFF), and atomic charges were calculated at the B3LYP/cc-pVTZ level. The total electrostatic charge of each system was neutralized by the addition of physiological concentrations of counterions. The time-step for integrating the equation of motion was 2 fs. Simulations were performed using periodic boundary conditions (PBC) in the NPT ensemble, i.e. at constant pressure (1 atm) and temperature (310 K), using a Langevin piston with a constant decay of 100 ps^−1^ and an oscillation period of 200 fs and a Langevin thermostat with a damping constant of 5 ps^−1^. Flexible unit cell was used with constant ratio in the x-y plane. Long-range electrostatic interactions were computed using the Particle Mesh Ewald method, with a fourth-order spline and 1 Å grid spacing. The POPC/water system was simulated for 30 ns to equilibrate, and then the following simulations were performed: three runs of spontaneous *trans* Ziapin2 insertion in the membrane (two for 175 ns and one for 300 ns); two runs of a single Ziapin2 in the membrane, one in *trans* and one in *cis* conformation (200 ns each); two runs each with four copies of Ziapin2 in the membrane, in *trans* and *cis* conformation (200 ns each); two runs each with eight copies of Ziapin2 in the membrane, in *trans* and *cis* conformation (200 ns each). When multiple copies of Ziapin2 were considered, they were distributing symmetrically in the upper and in the lower leaflet, aligned with the respective lipid headgroups. For the *cis* systems, the initial positions were determined by aligning the pyridine branches with those of the *trans* molecules. Two 200 ns simulations of four copies of Ziapin1 in the membrane were also performed, one with *trans* and one with *cis* conformations.

To determine the free energy profile for moving Ziapin2 from bulk water into the POPC bilayer we integrated a set of mean force values (i.e. minus the derivative of the free energy) calculated at 34 different positions of the molecule along the axis normal to the bilayer. The different values of the mean force were computed by restraining the center of mass of the pyridine nitrogen atoms of Ziapin2 at positions spaced by 1 Å along the normal with a force constant of 100 kcal·mol^−1^·Å^−2^. The center of mass of the bilayer was kept fixed in all mean force simulations. At each position the simulation lasted for 10-20 ns, until convergence of the mean force estimator was observed.

The raft model was generated as a 1:1:1 mixture of POPC, palmitoyl-sphingomyelin (PSM) and cholesterol (27 molecules of each component per leaflet) (49). The starting conformation for the raft in water was generated using pre-equilibrated lipid structures from the CHARMM-GUI webserver and then simulated for 50 ns to equilibrate in the new environment. Two systems were generated by inserting eight *trans* and *cis* Ziapin2 molecules, respectively, symmetrically distributed in the upper and in the lower leaflet with the pyridine rings aligned with the phosphate headgroups of POPC and PSM. Both systems were simulated for 200 ns.

Temperature Accelerated Molecular Dynamics (TAMD; 50) was used to accelerate the encounter of two *trans* Ziapin2 molecules in raft. TAMD is a method to enhance rare events sampling of a set of collective variables. The acceleration is obtained by employing extra dynamical variables tethered to the collective variables, which are evolved concurrently with the simulated system but at a higher temperature. By means of an effective adiabatic decoupling, the extra variables sample the Boltzmann equilibrium distribution at the higher temperature, while the rest of the system samples the same distribution at its lower temperature. Here, we accelerated the distance between centers of mass of two benzene rings in opposite Ziapin2 molecules. The force constant tethering the extra particle was 100 kcal·mol^−1^·Å^−2^, the friction coefficient of the particle dynamics was 100 ps·kcal·mol^−1^·Å^−2^, and the effective temperature 3000 K.

### Primary neuron preparations

Primary cultures of hippocampal neurons were prepared from embryonic 18-day C-57BL6/J mouse embryos (Charles River). Mice were sacrificed by CO_2_ inhalation, and embryos removed immediately by cesarean section. Briefly, hippocampi were dissociated by a 30-min incubation with 0.25 % trypsin at 37 °C and cells were plated on glass coverslips treated with Poly-L-lysine (0.1 mg/ml in borate buffer) in Neurobasal supplemented 2 mM L-glutamine, 2% B27, 100 μg/ml penicillin and 100 μg/ml streptomycin (Life Technologies). All animal manipulations and procedures were performed in accordance with the guidelines established by the European Community Council (Directive 2010/63/EU of March 4th 2014) and were approved by the Institutional Ethics Committee and by the Italian Ministry of Health. Ziapin2 diluted in DMSO at a concentration of 2 mM was added to the primary neuron preparation to reach final concentration ranging between 5 and 25 μM (final DMSO concentration < 1.5%). After an incubation time of 7 min at 37 °C, cells treated with vehicle or the photochromic molecule were washed with fresh solution and the coverslips were mounted on the imaging/electrophysiology chamber loaded with recording buffer.

### Fluorescence imaging of the plasma membrane

Live neurons were loaded with CellMask™ Deep Red plasma membrane stain (1 μl/ml) or Vybrant™ Alexa Fluor™ 555 lipid rafts labeling kit (ThermoFisher) for the evaluation of the localization of Ziapin2 at the plasma membrane and lipid rafts, respectively, and Hoechst 33342 (1 μM) for nuclear visualization. Briefly, primary neurons were exposed to 5 μM Ziapin2 for 7 min; the molecule was then washed away and neurons were treated with either CellMask™ Deep Red or Vybrant™ Alexa Fluor™ 555, following the manufacturer instructions. Coverslips with cells were washed with fresh medium, and mounted on a confocal laser-scanning microscope (CLSM) Leica SP8 (Leica Microsystem) for live-cell z-series stack acquisition of consecutive confocal sections. Fluorescence levels and co-localization analysis were carried out using the Fiji software and Coloc2 plugin (ImageJ).

### Electrophysiology

Whole-cell patch-clamp recordings of hippocampal neurons (between 14 and 18 day in vitro, DIV) were performed at room temperature using borosilicate patch pipettes (3.5-5.0 MΩ) and under GΩ patch seal. The extracellular solution contained (in mM): 135 NaCl, 5.4 KCl, 1 MgCl_2_, 1.8 CaCl_2_, 5 HEPES, 10 glucose adjusted to pH 7.4 with NaOH. The intracellular solutions used for neuron recordings contained (in mM): neurons: 126 K-Gluconate, 4 NaCl, 1 MgSO_4_, 0.02 CaCl_2_, 0.1 EGTA, 10 Glucose, 5 Hepes, 3 ATP-Na_2_, and 0.1 GTP-Na. To block excitatory and inhibitory synaptic transmission, recordings were performed in the presence of D-AP5 (50 μM), CNQX (10 μM), and bicuculline (30 μM) (Tocris). Recordings were carried out using either an Axopatch 200B (Molecular Devices) or an EPC10 (HEKA Elektroniks) amplifier. Responses were amplified, digitized at 10 or 20 kHz and stored with either Clampfit (Molecular Devices) or Patchmaster V2.73 (HEKA Elektroniks). pCLAMP 10 and FitMaster v2×90.1 were employed for data analysis, together with Prism 6.07 (GraphPad) and OriginPro 9 (OriginLab).

### Photostimulation

Illumination of neurons during electrophysiological experiments was provided by an LED system (Lumencor Spectra X) fibre-coupled to an upright Nikon FN1 microscope. The light source emission peaked at 470 nm to match the Ziapin2 absorption spectrum and the power density ranged from 18 to 50 mW/mm^2^, as measured at the output of the microscope objective.

### In vivo experiments

*Surgery*. In vivo optical stimulations were performed in C57BL6 mice that had been previously injected with either DMSO (N=3) or Ziapin2 (N=3). Mice were anaesthetized with Isoflurane and placed in a stereotaxic frame, where anesthesia was maintained with an isoflurane flow. Ziapin2 (200 μM in 1 μl 10% DMSO in PBS) or vehicle was injected with a 5 μl Hamilton syringe in the primary somatosensory cortex (S1ShNc) of the left hemisphere using the following stereotactic coordinates: 2 mm anterior to lambda, 2 mm lateral to midline, and – 723 μm ventral to brain surface. Injection was done at a rate of 100 nl/min with a nano-jector (World Precision Instruments, Fl, USA). After a 5 min wait for the molecules to diffuse within the brain, the craniotomy was extended by 1 mm laterally and 1 mm medially to accommodate the microwire array (16 electrodes in 2 rows of 8, 33 μm diameter, 250 μm pitch, 375 μm between rows (Tucker Davis Technologies). The two central microwires were inserted at a depth of 723 μm, before being cleaned with saline and topped with silicone sealant (Kwik-cast, WPI). A hole was drilled 1 mm caudally to the injection hole and a fiber optic cannula (MFC_400/430-0.66_10mm_ZF1.25_FLT, Doric Lenses) was inserted at a depth of 1 mm and an angle of 65° from the vertical. Two surgical screws were inserted in the skull contralateral to the implants to be used as reference/ground for the microwires and support. The skull was then covered with dental cement and Diclofenac was systemically administered at a dose of 100 μl/ 20g. Mice were left to recover for 30 min before electrophysiological recordings. *In vivo recordings*. Light stimulation was delivered at 0.25 Hz with 40 % jitter for either 20 or 200 ms at irradiances of 4, 25, 43 or 160 mW/mm^2^ with a 473 nm laser (Shanghai Dream Lasers) to the freely moving rats. Each condition was repeated 25 times. Extracellular signals in response to stimulation were amplified, digitized and sampled at 1017 Hz by commercially available hardware (System 3, Tucker-Davis Technologies) before being saved for offline analysis using custom Matlab scripts (The Mathworks). During acquisition, data were high-pass (1 Hz) and low pass (100 Hz) filtered to extract local field potentials (LFPs).

### Statistical analysis

Data were expressed as means ± SEM for number of cells or preparations (N). Normal distribution was assessed using D’Agostino-Pearson’s normality test. To compare two sample groups, Student’s *t*-test or Mann-Whitney *U*-test was used. To compare more than two sample groups, ANOVA followed by Holm-Sidak’s post-hoc test or Kruskal-Wallis test followed by the Dunn’s post-hoc test were used. Statistical analysis was carried out using OriginPro-8 (OriginLab) and Prism (GraphPad).

