## Supplementary Materials for "A Membrane-Targeted Photoswitch Potently Modulates Neuronal Firing"

### SUPPLEMENTARY TEXT

#### Ziapin 2 closely overlaps with the plasma membrane reporter Cell Mask

Given the substantial overlap between the emission spectrum of Ziapin2 ($\lambda_{ex}$ 488 nm; $\lambda_{em}$ 566-620 nm) and the excitation spectrum of Cell Mask ($\lambda_{ex}$ 633 nm; $\lambda_{em}$ 650-750 nm), we addressed the possibility that, in the presence of a strict co-localization, Ziapin2 emission could excite Cell Mask (**Fig. S3a-c)**. To ascertain this, primary neurons exposed to either DMSO or Ziapin2 were live-stained with Cell Mask and imaged by confocal microscopy by exciting Ziapin2 at 488 nm and detecting fluorescence emission alternatively at 566-620 nm ($\lambda_{em}$ of Ziapin2) and 650-750 nm ($\lambda_{em}$ of Cell Mask). While no signal was detected in the absence of Ziapin2, an intense signal in the 650-750 nm range was detected when neurons were pre-exposed to Ziapin2 (**Fig. S3a-c**). The resulting morphological pattern was closely similar to that obtained by directly exciting Cell Mask at 633 nm (**Fig. S3a**). These data indicate a close overlap between Ziapin2 and the live plasma membrane reporter Cell Mask, further demonstrating that the molecule is strictly targeted to the membrane bilayer.

#### The reversible membrane thinning due to Ziapin2 photoisomerization also occurs in lipid rafts

To verify whether the same membrane push-pull effect also occurs in lipid rafts, we adopted a well-accepted raft model. Due to the increased rigidity and thickness of the raft environment that makes the time-scale of the encounter between opposing Ziapin2 molecules longer than the length of our simulated trajectory (200 ns), we accelerated the encounter process by employing the TAMD-enhanced sampling method (see Online Methods; 50). Specifically, we accelerated the motion of a variable describing the distance between the centers of mass of the benzene rings in two opposing Ziapin2 molecules, and observed that the two molecules came together within 20 ns, forming an arrangement similar to the one that spontaneously occurred in the membrane simulation. Similar to what observed in the POPC membrane, the interaction between the two Ziapin2 molecules caused a local depression in the raft, as a consequence of the inward displacement of the POPC and PSM phosphate atoms coordinating the Ziapin2 pyridine rings. Then, starting from the final conformation of the TAMD trajectory, we ran 100 ns standard simulation to check the stability of the obtained conformation. During this trajectory, the arrangement of the two interacting Ziapin2 molecules and the associated pinching of the raft were stably maintained (**Fig. S5**). Thus, the same membrane thinning mechanism resulting from the interaction of opposing *trans* Ziapin2 molecules is observed also within the preferred membrane environment of lipid rafts.

**Model of membrane capacitance changes induced by Ziapin2**

The plasma membrane can be assimilated to a planar capacitor whose capacitance is:

$$C = k_d \varepsilon_0 \frac{A}{l}$$

A change in capacitance at fixed charge, "open circuit" condition (i.e., before the induced current reestablishes the equilibrium) induces a change in membrane voltage. The change in capacitance can be due to a change in membrane dielectric constant, membrane thickness or membrane area, according to:

$$dC = dk_d \varepsilon_0 \frac{A}{l} - k_d \varepsilon_0 \frac{A}{l^2} dl + k_d \varepsilon_0 \frac{dA}{l}$$

Assuming that the change in area (dA) is negligible, two contributions describe the variation induced in capacitance by the insertion of the photochromic molecule, namely a change in dielectric constant, $dk_d$, and/or a change in thickness, *dl*.

The average initial capacitance of the cell in the resting state and under control conditions is $C_0$ = 20-30 pF. In the presence of the photochromic molecule in the medium, it becomes $C_d$ = 45-55 pF. Taking into account the propagation of experimental errors, the relative increase $\frac{dC}{C_0}$ is $\approx 100\%$. If this were due to thickness modification only $\frac{dC}{Co} = -\frac{dl}{l}$. The estimated percent change in thickness due to the dimerization of the chromophores, according to our simulations, is $\frac{\Delta l}{l} \leq 20\%$ that is definitely smaller than the observed change in capacitance. Thus, it is likely that the simulations underestimate the thinning effect due to the limited number of molecules that can be considered with respect to the real situation. In addition, the capacitance jump upon the initial addition of the compound in the dark could be due to the combination of thickness change and a change in the membrane dielectric constant. Upon removal of the unbound compound and subsequent illumination, the observed variation of capacitance (about 10%) is fully consistent with the extent of photoisomerization-induced membrane relaxation predicted by the simulations.

### SUPPLEMENTARY FIGURES

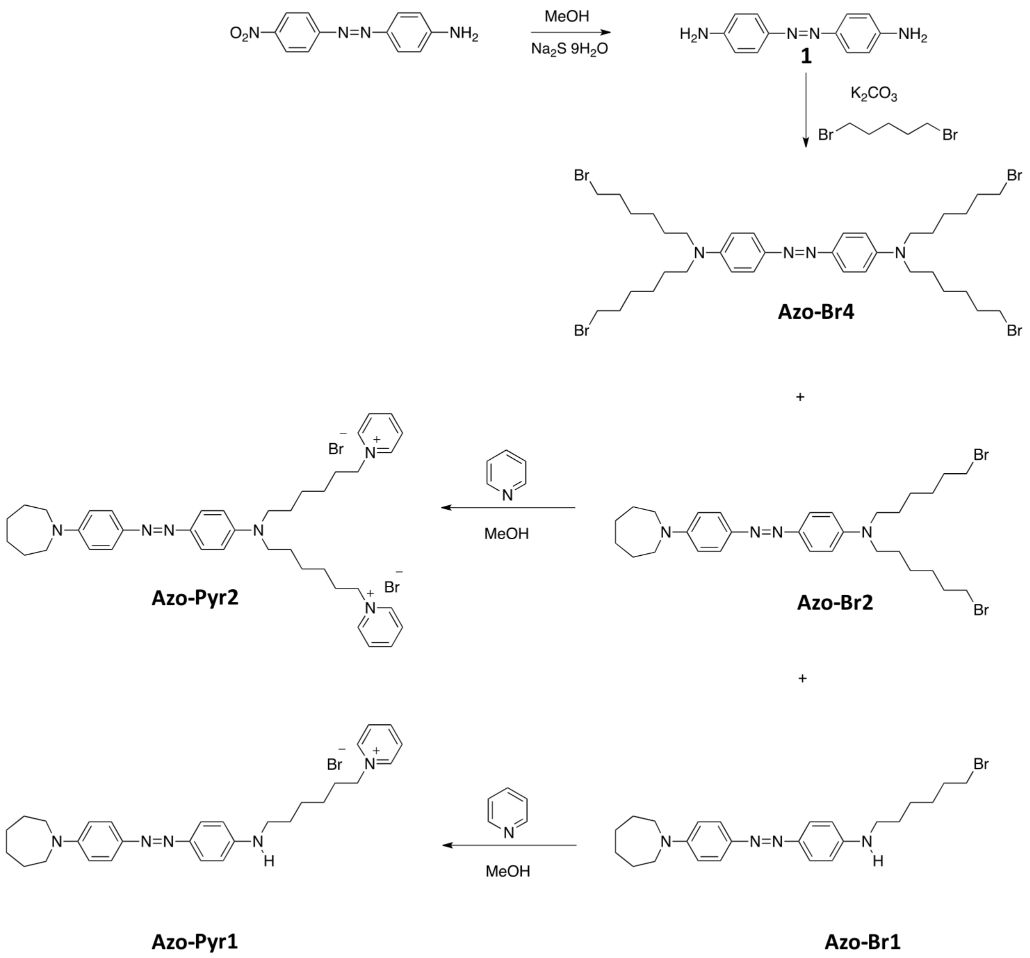


**Figure S1. Synthetic route of the photochromic azobenzenes Ziapin1 (Azo-Pyr1) and Ziapin2 (Azo-Pyr2)**
For further details, see Methods.

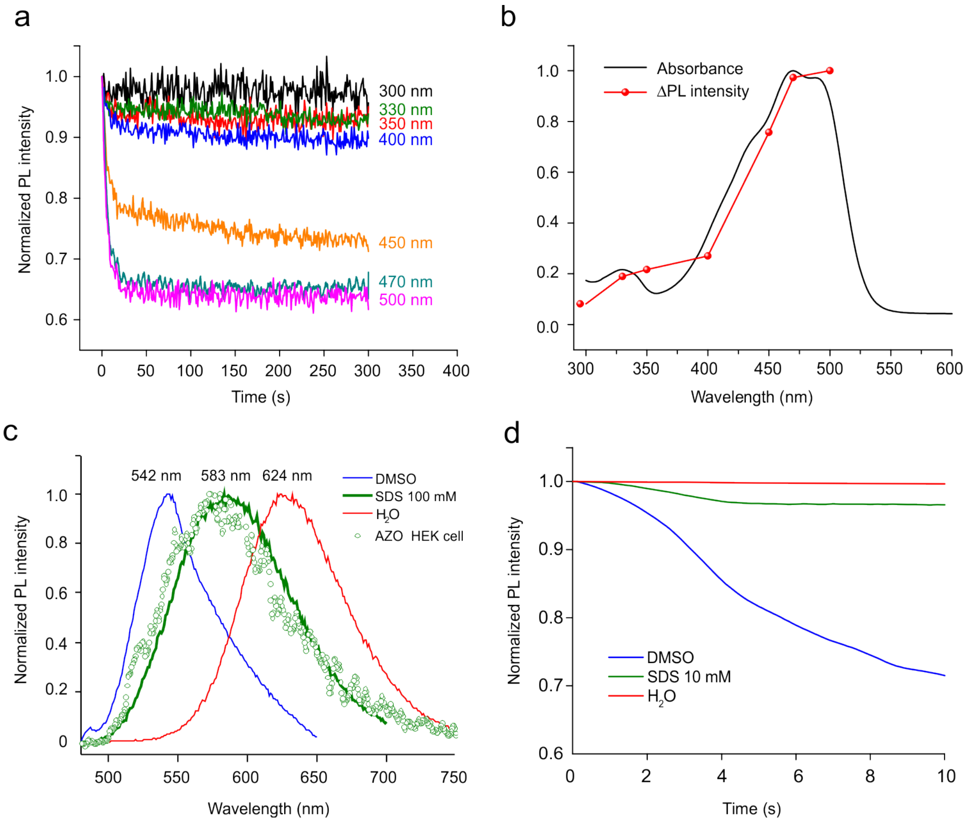


**Figure S2. Fluorescence emission of Ziapin2 in mixed environments**

**a.** Time-dependent emission of Ziapin2 (25 μM) in DMSO at 540 nm under illumination at various wavelengths.

**b.** Profiles of normalized fluorescence and absorption spectrum as a function of the excitation wavelength of Ziapin2 (25 μM) in DMSO.

**c.** Emission spectra of Ziapin2 in DMSO (100%), SDS (100 mM, above the critical micellar concentration of 8.2 mM @ 25 °C), HEK293 cells and water. The fluorescence emission curve of Ziapin2 in SDS lies exactly in between those in DMSO (peak at 542 nm) and water (peak at 624 nm). Remarkably, the fluorescence spectrum taken from the Ziapin2-HEK293 cells largely overlaps with the emission from the Ziapin2-SDS sample, suggesting that SDS micelles mimic the local environment of the cell membrane and that Ziapin2 molecules in cells may be mostly trapped in the membrane. Under the SDS and HEK293 cell conditions, Ziapin2 is expected to be partly solvated by DMSO, partly trapped in the hydrophobic membrane/micelle environment, and partly aggregated in water. Accordingly, the measured spectrum is wider than those in pure DMSO or water.

**d.** Photoluminescence dynamics of Ziapin2 in DMSO, SDS (100 mM) and water acquired by illuminating with a Xenon lamp (450 nm) and collecting the emission at 540 nm, 580 nm and 620 nm for respectively.

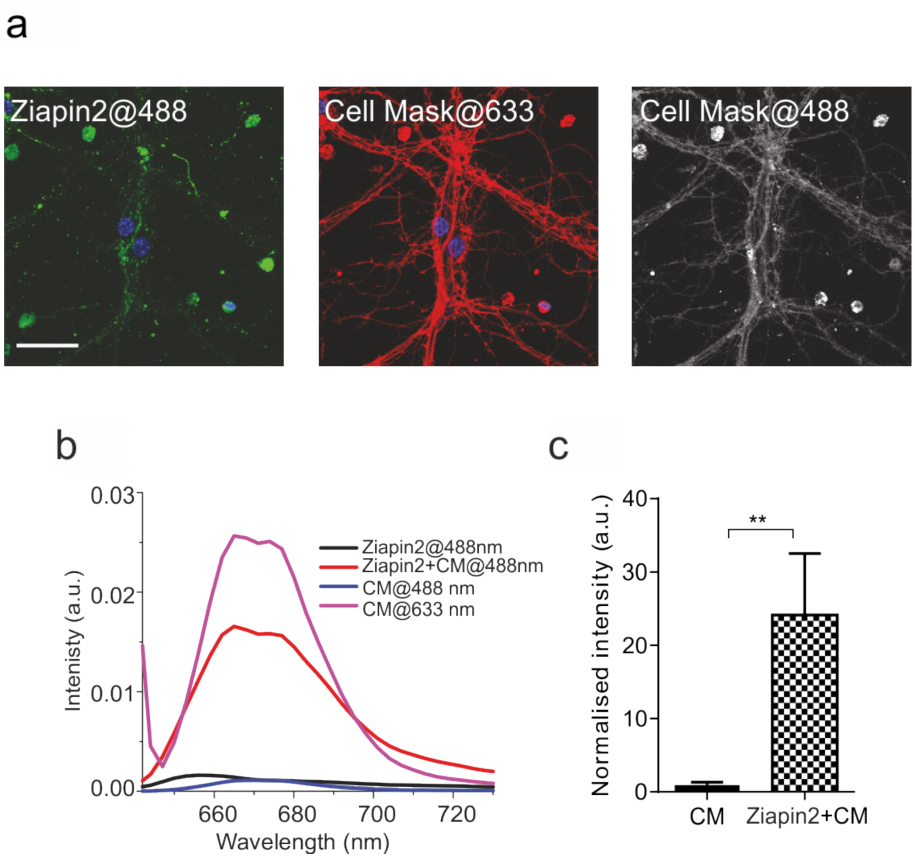


**Figure S3. Codistribution of Ziapin2 with the plasma membrane marker Cell Mask**

**(a)** Primary neurons exposed to Ziapin2 (green at $\lambda_{ex}$ 488 nm) for 7 min were stained with Cell Mask (red at $\lambda_{ex}$ 633 nm). From left to right, the following conditions are shown: Ziapin2 imaged by standard excitation at 488 nm and emission at 566-620 nm; Cell Mask imaged by standard excitation at 633 and emission at 650-750 nm; sample excited at 488 nm (specific for the Ziapin2) and imaged at 650-750 nm (specific for Cell Mask). The pattern of the last acquisition was identical to the Cell Mask pattern, suggesting that Ziapin2 was located in the plasma membrane to excite Cell Mask. Scale bar, 20 $\mu$m.

**(b,c)** This phenomenon was quantified (**b**) and compared to fluorescent intensity signal of Cell Mask excited at 488 nm and detected at 650-750 nm in absence of Ziapin2 molecules (**c**). Data are expressed as mean ± SEM; ** $p<0.01$ (n = 30 cells for each condition from 3 independent animal preparations).

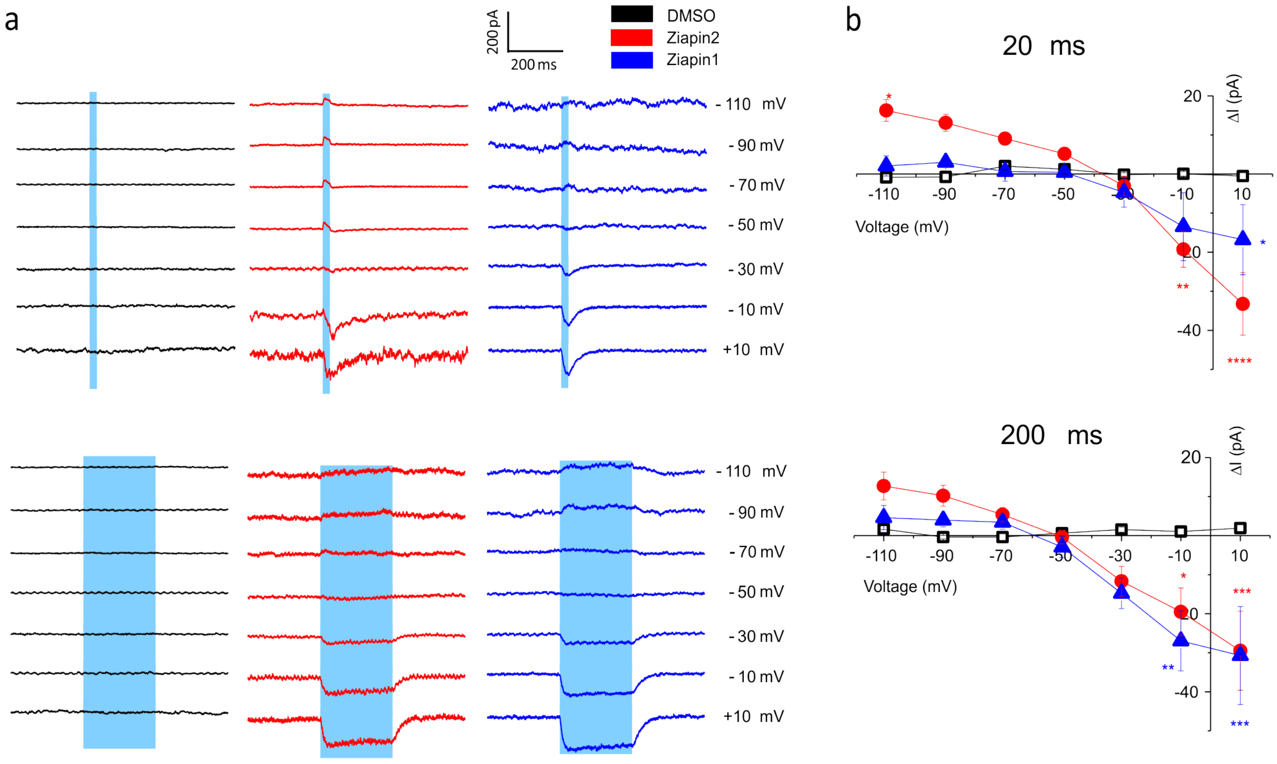


**Figure S4. Light-dependent Current/Voltage relationships in neurons loaded with Ziapin2 or Ziapin1**

(**a**) Representative averaged traces recorded in voltage-clamp configuration from neurons loaded with either DMSO (0.25% v/v, black traces), Ziapin2 (5 µM in DMSO, red traces) or Ziapin1 (5 µM in DMSO, blue traces). Neurons clamped at increasing voltages from -110 to 10 mV were photostimulated with 20 ms (upper panels) or 200 ms (lower panels) light pulses (470 nm; 18 $mW/mm^2$; cyan-shaded areas).

(**b**) Current/voltage relationships in response to illumination with 20 (upper panel) or 200 (lower panel) ms pulses built from the experiments depicted in (**a**) from neurons loaded with DMSO (black symbols), Ziapin2 (red symbols) or Ziapin1 (blue symbols). * p<0.05; ** p<0.01; *** p<0.001; Friedman's two-way ANOVA/Dunn's tests. N=9 (DMSO), N=22 (Ziapin2), N=11 (Ziapin1).

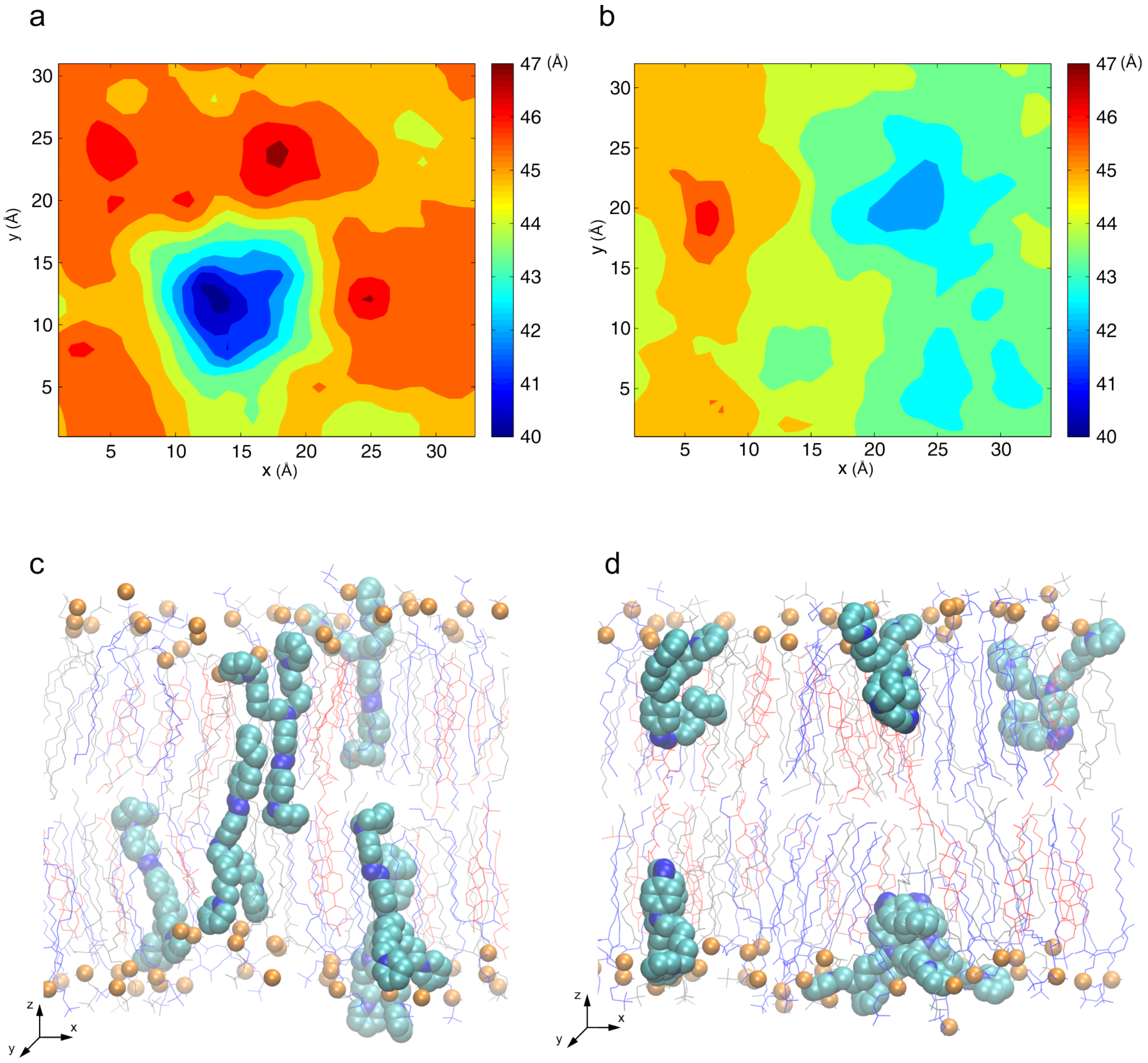


**Figure S5. Molecular Dynamics simulations of Ziapin2 in a raft membrane model**

(**a,b**) Average membrane thickness maps, shown perpendicular to the bilayer plane, for the high-concentration (8 molecules) simulations of Ziapin2 in *trans* (**a**) and *cis* (**b**) conformation, respectively. Axis and thickness values are in Å.

(**c,d**) Snapshots from the simulations of eight Ziapin2 molecules in *trans* (**c**) and *cis* (**d**) conformation and embedded in a raft membrane model (1:1:1 mixture of POPC, palmitoyl-sphingomyelin, PSM, and cholesterol). POPC lipid molecules are shown as grey lines, and phosphate atoms as orange spheres; cholesterol and PSM molecules are shown as red and blue lines, respectively; water molecules and ions are not reported for clarity.

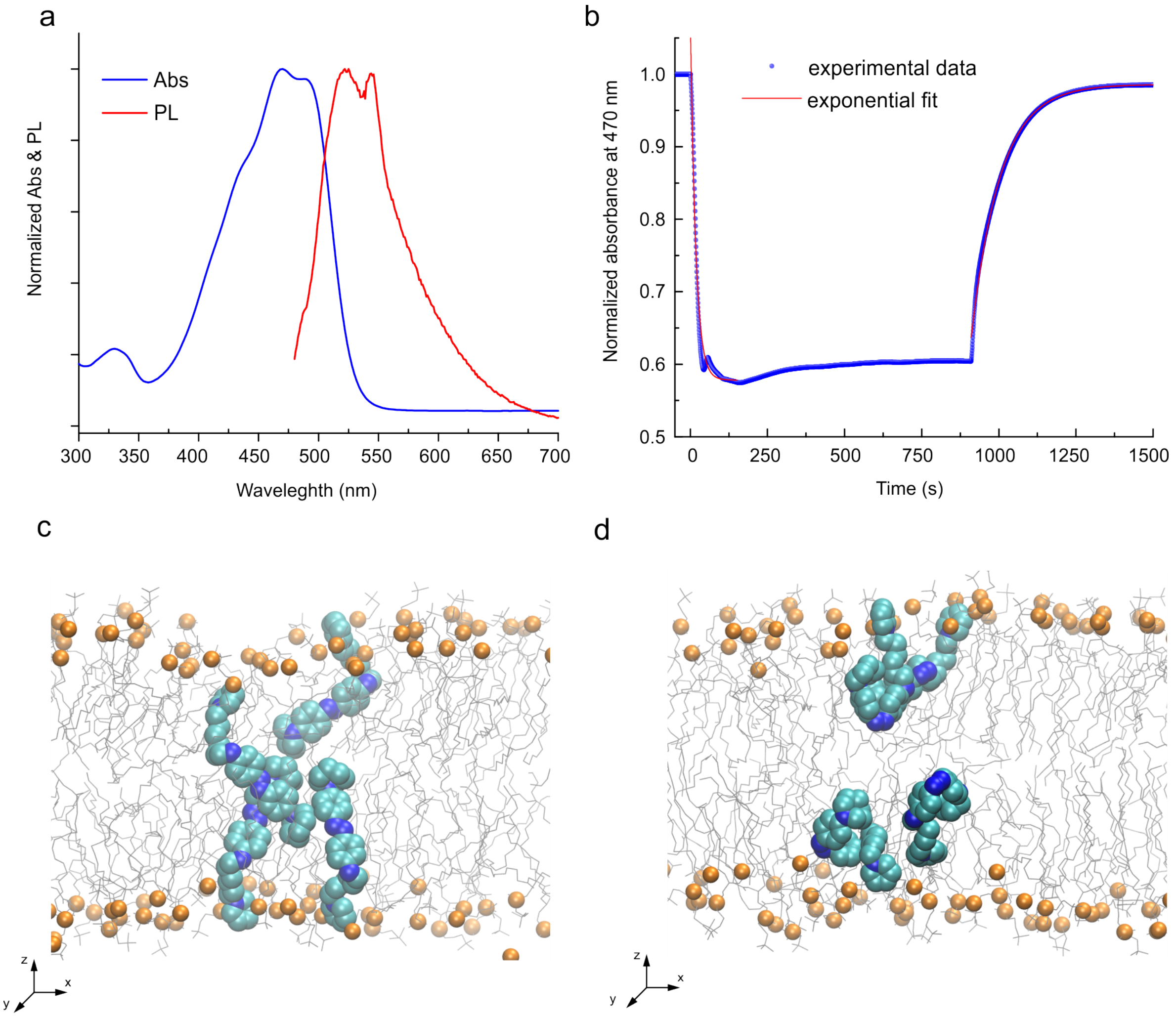


**Figure S6. MD simulations in a membrane model and spectroscopic characterization of Ziapin1**
(**a,b**) Absorption (blue) and PL (red) spectra of Ziapin1 (25 µM) in DMSO (**a**) and time-course of the absorbance peak at 470 nm upon illumination with a diode laser at 450 nm (**b**). Note that Ziapin1 reaches a photostationary state in ≈ 100 s, similar to Ziapin2, while the $t_{1/2}$ of recovery (64 s) is slightly shorter than that of Ziapin2 (108 s).
(**c,d**) Snapshot extracted from the simulation of four Ziapin1 molecules in *trans* (**c**) and *cis* (**d**) conformation embedded in a membrane model (POPC lipid). Lipid molecules are shown as grey lines, and phosphate atoms as orange spheres; water molecules and ions are not reported.

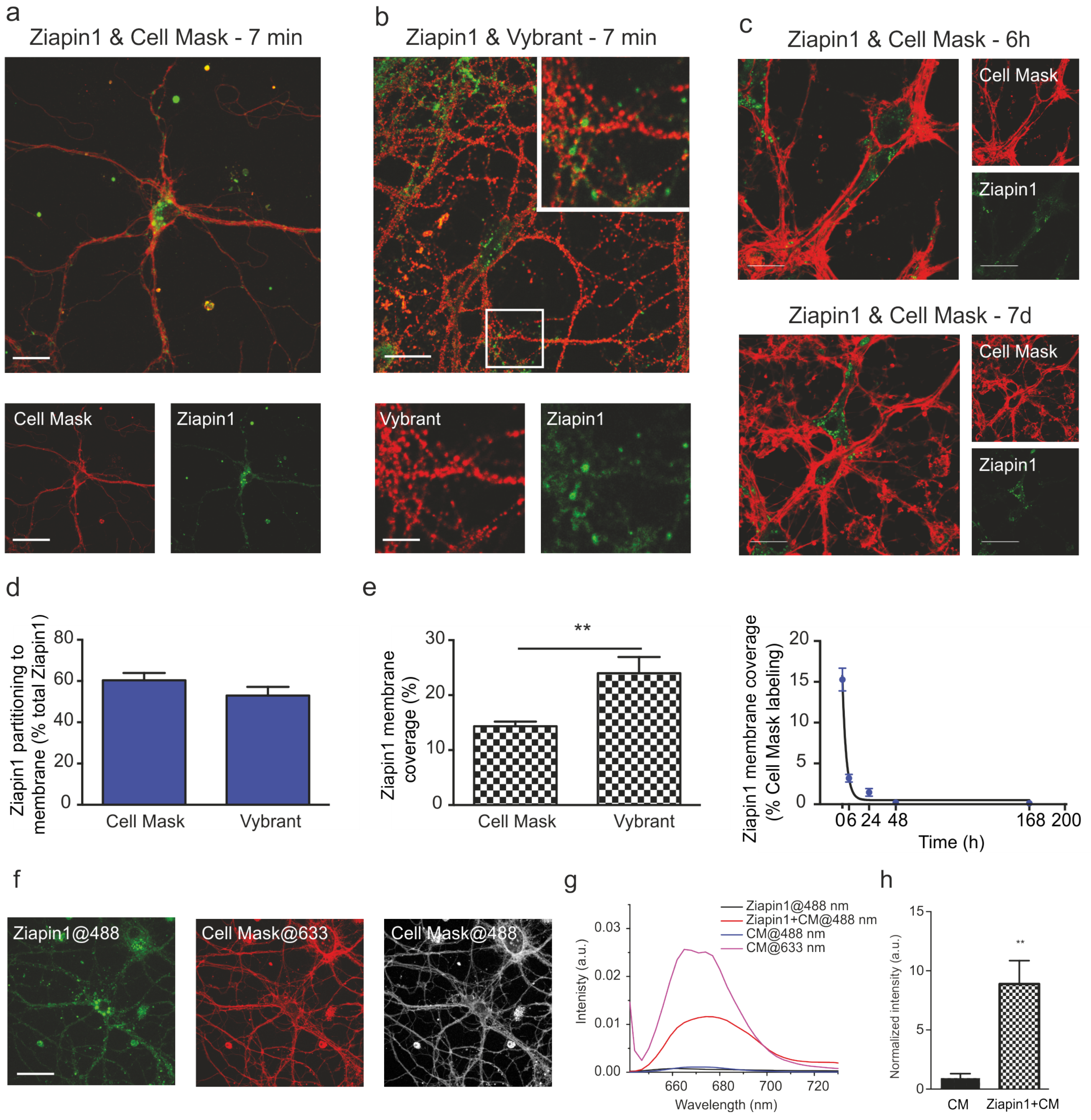


**Figure S7. Targeting of Ziapin1 to the neuronal plasma membrane**

(**a,b**) Primary neurons exposed to Ziapin1 for 7 min were stained with specific membrane dyes to evaluate and quantify the co-localization of the molecule with the plasma membrane. Cell Mask staining was used to visualize the whole membrane of neurons (**a**, in red; Ziapin1 in green), while Vybrant™ Alexa Fluor™ 555 lipid rafts labeling kit was used for the identification of lipid rafts (**b**, in red; Ziapin1 in green). Scale bars: 20 and 40 μm for large and small panels, respectively (**a**); 20 and 6 μm for large and small panels, respectively (**b**).

(**c**) The Cell Mask/Ziapin1 double staining is shown 6 h (upper panel) and 7 d (lower panel) after the initial exposure. Scale bars: 10 and 20 μm for large and small panels, respectively (6h); 20 and 40 μm for large and small panels, respectively (7 d).

(**d**) After Cell Mask/Ziapin1 and Vybrant/Ziapin1 double staining and z-stack confocal imaging, the amount of Ziapin1 co-localizing with the plasma membrane and with lipid rafts was quantified as the percentage of molecule present on the respective membrane staining over the total cell Ziapin1 after 7 min of incubation and washout (time 0).

(**e**) Percentage of total plasma membrane/lipid rafts staining covered by Ziapin1 after 7 min of incubation and washout (time 0; left panel). On the right, the Ziapin1 coverage of the plasma membrane is shown as a function of time up to 7 days after the initial exposure. The progressive decrease of Ziapin1 membrane coverage was fitted using a one-component exponential decay function ($t_{1/2}$ = 2.4 h).

(**f**) From left to right, the following conditions are shown: Ziapin1 imaged by standard excitation at 488 nm and emission at 566-620 nm; Cell Mask imaged by standard excitation at 633 and emission at 650-750 nm; sample excited at 488 nm (specific for the Ziapin1) and imaged at 650-750 nm (specific for Cell Mask). The pattern of the last acquisition is identical to the Cell Mask pattern, suggesting that Ziapin1 was located in the plasma membrane to excite Cell Mask. Scale bar, 20 μm.

(**g,h**) This phenomenon was quantified (**g**) and compared to fluorescent intensity of Cell Mask excited at 488 nm and detected at 650-750 nm in absence of Ziapin1 molecules (**h**). Data are expressed as means ± SEM; ** $p < 0.01$ (n = 30 cells for each condition from 3 independent animal preparations).

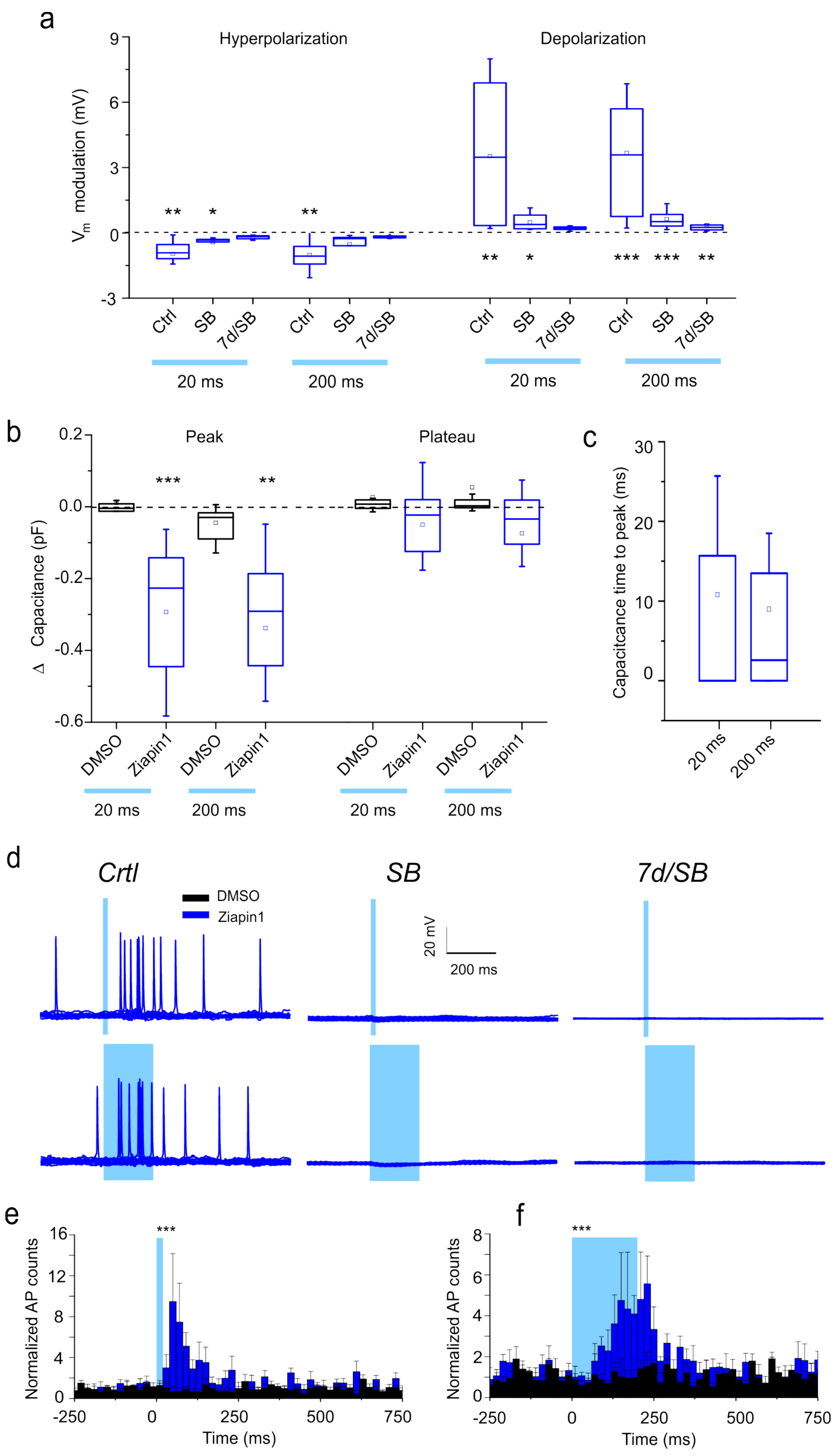
a
Hyperpolarization
Depolarization
$V_m$ modulation (mV)
9
6
3
0
-3
** * **
** * *** *** **
Ctrl
SB
7d/SB
20 ms
200 ms
b
Peak
Plateau
Δ Capacitance (pF)
0.2
0.0
-0.2
-0.4
-0.6
*** **
DMSO
Ziapin1
20 ms
200 ms
c
Capacitcance time to peak (ms)
30
20
10
0
20 ms
200 ms
d
Crtl
SB
7d/SB
DMSO
Ziapin1
20 mV
200 ms
e
f
Normalized AP counts
16
12
8
4
0
***
-250
0
250
500
750
Time (ms)

**Figure S8. Effects of Ziapin1 on the passive and active membrane properties in primary hippocampal neurons**

Primary neurons were loaded with either DMSO (see Fig. 5) or Ziapin1 (5 µM in DMSO, blue traces) and subsequently recorded in the presence (SB) or absence (Ctrl) of synaptic blockers in response to 20 or 200 ms light stimulation (470 nm; 18 $mW/mm^2$). Neurons were also recorded 7 days after loading in the presence of synaptic blockers (7d/SB).

(**a**) Box plots of the peak hyperpolarization (left) and peak depolarization (right) changes in response to 20/200 ms light stimulation. Hyperpolarization and depolarization were measured as the minimum and maximum voltage, respectively, reached within 350 ms from the light-onset. * p<0.05; ** p<0.01; *** p<0.001; Ziapin1 *vs* DMSO, Mann Whitney *U*-test. (Ziapin1 - Ctrl, SB, 7d/SB: N = 15, 15, 19).

(**b,c**) Box plots of the peak (left) and plateau (right) capacitance changes (**b**) and time to peak capacitance change (**c**) evoked in neurons exposed to either DMSO or Ziapin1 and stimulated by 20 and 200 ms light stimuli in the presence of synaptic blockers. (Ziapin1, N = 12; DMSO, N=9). ** p<0.01; *** p<0.001, Ziapin1 *vs* DMSO, Mann-Whitney *U*-test. ° p<0.05, 200 *vs* 20 ms, one-way ANOVA/Holm-Sidak's tests.

(**d**) Representative current-clamp recordings of AP firing in primary neurons loaded with Ziapin1 (5 µM in DMSO, blue traces) in the absence (Ctrl) or presence of synaptic blockers (SB). Neurons were also recorded 7 days after loading in the presence of synaptic blockers (7d/SB). Neurons were challenged with either 20 ms (upper panel) or 200 ms (lower panel) light stimulation (470 nm; 18 $mW/mm^2$) shown as a cyan-shaded area.

(**e,f**) Light-induced firing was analyzed by building peristimulus time histograms (PSTHs; bin=20 ms) from the firing rate of neurons loaded with either DMSO (black bars) or Ziapin1 (blue bars) recorded in the absence of synaptic blockers in response to 20 ms (**e**) or 200 ms (**f**) light stimulation. (Ziapin1 - Ctrl: N=7 and N=5 for 20 and 200 ms respectively; DMSO - Ctrl: N=4). *** p<0.001; Ziapin1 *vs* DMSO, Mann-Whitney *U*-test.
